# Systematic analysis of the effects of splicing on the diversity of post-translational modifications in protein isoforms using PTM-POSE

**DOI:** 10.1101/2024.01.10.575062

**Authors:** Sam Crowl, Maeve Bella Coleman, Andrew Chaphiv, Ben T. Jordan, Kristen M. Naegle

**Affiliations:** University of Virginia, Department of Biomedical Engineering and the Center for Public Health Genomics, Charlottesville, VA, 22903

## Abstract

Post-translational modifications (PTMs) and splicing are important regulatory processes for controlling protein function and activity. Despite examples of interplay between alternative splicing and cell signaling in literature, there have been few detailed analyses of the impacts of alternative splicing on PTMs, partly due to difficulties in extracting PTM information from splicing measurements. We developed a computational pipeline, PTM Projection Onto Splice Events (PTM-POSE), to identify “prospective” PTM sites in alternative isoforms and splice events recorded in databases using only the genomic coordinates of a splice event or isoform of interest. Importantly, PTM-POSE integrates various PTM-specific databases and tools to allow for deeper analysis of the individual and global impact of spliced PTMs on isoform function, protein interactions, and regulation by enzymes like kinases. Using PTM-POSE, we performed a systematic analysis of PTM diversification across isoforms annotated in the Ensembl database. We found that 32% of PTMs are excluded from at least one Ensembl isoform, with palmitoylation being most likely to be excluded (49%) and glycosylation and crotonylation exhibiting the highest constitutive rates (75% and 94%, respectively). Further, approximately 2% of prospective PTM sites exhibited altered regulatory sequences surrounding the modification site, suggesting that regulatory or binding interactions might be different in these proteoforms. When comparing splicing of phosphorylation sites to measured phosphorylation abundance in KRAS-expressing lung cells, differential inclusion of phosphorylation sites correlated with phosphorylation levels, particularly for larger changes in inclusion (*>* 20%). To better understand how splicing diversification of PTMs may alter protein function and regulatory networks in specific biological contexts, we applied PTM-POSE to exon utilization measurements from TCGASpliceSeq of prostate tumor samples from The Cancer Genome Atlas (TCGA) and identified 1,489 PTMs impacted by ESRP1-correlated splicing, a splicing factor associated with worsened prognosis. We identified protein interaction and regulatory networks that may be rewired as a result of differential inclusion of PTM sites in ribosomal and cytoskeletal proteins. We also found instances in which ESRP1-mediated splicing impacted PTMs by altering flanking residues surrounding specific phosphorylation sites that may be targets of 14-3-3 proteins and SH2 domains. In addition, SGK1 signaling was found to be influenced by ESRP1 expression through increased inclusion of SGK1 substrates in ESRP1-expressing patients. Based on validation in a separate prostate cancer cohort from the Chinese Prostate Cancer Genome and EpiGenome Atlas (CPGEA), this correlated with increased phosphorylation of SGK1 substrates, particularly when SGK1 was predicted to be active. From this work, we highlighted the extensive splicing-control of PTM sites across the transcriptome and the novel information that can be gained through inclusion of PTMs in the analysis of alternative splicing. Importantly, we have provided a publicly available python package (PTM-POSE: https://github.com/NaegleLab/PTM-POSE) and all associated data for use by the broader scientific community to allow for continued exploration of the relationship between splicing and PTMs.

## Introduction

While protein function is often treated as a static entity for multi-omic analyses, proteins can take many different ‘proteoforms’ with distinct functions through alternative splicing, post-translational modifications (PTMs), and other mechanisms [1]. While nearly all multi-exonic genes undergo alternative splicing to produce multiple unique protein isoforms, the functional significance of many splice variants remains poorly understood [2, 3]. Among the many factors that dictate isoform function, gain or loss of post-translational modification (PTM) sites has been illustrated to have significant functional consequences, such as for a glycosylation site found in an isoform of neuroligin-1 that regulates its interaction with neurexin, altering synapse formation [4]. Previous research has suggested that PTM sites, particularly those found within disordered regions of proteins, are enriched within tissue-specific exons (differentially expressed across tissues) [5, 6], although some work disputes this idea [7]. Furthermore, alternatively spliced regions of proteins are enriched for linear motifs corresponding to several PTM-specific binding domains important for facilitating protein interaction and signaling networks, including SH2 and WW domains [8]. However, the relationship between splicing and PTMs remains underexplored, as most proteomic experimental pipelines cannot capture different protein isoforms from small peptide fragments. Hence, the majority of published proteomic experiments, which are then incorporated into PTM-databases, assign PTMs to a single reference protein, typically the “canonical” isoform [9]. Consequently, existing analyses of PTM diversity across isoforms rely on on a small subset of data (such as tissue-specific exons) [5–7], sparse UniProt annotations of PTMs [8, 10], and/or consider all PTMs as one entity, limiting the broad utility of these results. Additionally, no study has systematically explored how alternative splicing may modify the flanking sequence surrounding a PTM, which fundamentally encodes the enzymatic recognition that ‘wires’ PTMs to their regulatory enzymes and sometimes reader domains [12] [13].

In recent years, many databases and tools have been developed to facilitate exon- or isoform-specific functional analysis of experimentally measured splicing events. These tools provide exon- or isoform-specific functional annotations [14–16], protein interactions [17–20], and tissue and disease expression [21, 22], among other information. While some newer tools annotate splice events or isoforms with PTMs from either UniProt [29] or PhosphoSitePlus [31], many of these databases/tools cannot be easily applied to new splice events or datasets, instead focusing only on a specific subset of data (such as cancer and tissue specific splice events [21]) or reliant on previously annotated transcripts [18]. More importantly, they contain limited, if any, functional or regulatory information related to impacted PTM sites, resorting to only reporting that a PTM is impacted. This makes it difficult to assess the functional relevance of these PTM sites. For example, PTMs can trigger protein-interactions through PTM-specific recognition domains: bromodomains bind to acetyl lysines [23], WW domains and 14-3-3 proteins bind phosphorylated serines and threonines [24], and ubiquitin-binding domains bind ubiquitinated lysines [25] (Supplementary Figure 1). However, while tools like DIID [20], NEASE [15], and LINDA [19] predict impacts of splicing on protein interaction networks, they do not consider changes to inclusion or motifs of PTMs, relying instead on domain-domain interaction information. Further, none of these tools consider the flanking residues surrounding PTMs impacted by splice events that disrupt linear motifs required for certain protein interactions. Therefore, there is a need for new tools that enable more comprehensive, data-agnostic analyses of the relationship between splicing and PTMs that considers the diversity of PTMs, their potential functions, and the different ways in which splicing can impact PTMs in specific biological and experimental contexts.

Here, we developed PTM Projection Onto Splice Events (PTM-POSE), an open-source python-based method to identify “prospective” PTM sites associated with measured splice events or isoforms. PTM-POSE employs a data-agnostic approach requiring only the genomic coordinates of the isoform(s) or event(s) of interest, enabling broad applicability to any measured splice event or exon, such as those quantified from RNA-sequencing data. Further, we integrated a suite of downstream analysis modules to assess how splice-affected PTMs impact: 1) protein function, 2) protein interactions, and 3) enzyme regulation, including kinase-substrate relationships. These capabilities allow researchers to identify global patterns in PTM-related isoform function, such as coordinated changes to kinase-substrate availability resulting from differential exon inclusion. Further, PTM-POSE uniquely identifies PTM sites with altered flanking sequences between isoforms and evaluates how these changes may affect binding motifs and enzyme recognition. This allows for a more comprehensive analysis of the functional consequences of splicing on PTMs, as well as the potential for splicing to alter the regulatory landscape of proteins through changes to PTM sites.

To first establish the extent of PTM diversification through splicing, we applied PTM-POSE to alternative isoforms recorded in the Ensembl database. We found that splicing significantly influences the PTM landscape across the proteome, with *≈* 30% of PTMs excluded from at least one isoform. Certain modification types are more likely to be maintained across different isoforms, such as glycosylation. Further, we found that a subset of prospective PTM sites (*≈* 2%, amounting to 14,850 prospective PTM sites) exhibit altered flanking sequences that may serve to change the regulatory and binding interactions associated with the PTM. Importantly, changes in inclusion of phosphorylation sites correlated with their measured abundance by mass spectrometry in two different datasets, highlighting that measured splicing-level changes can impact PTM abundance. This analysis represents one of the most comprehensive examinations to date of how alternative splicing may reshape the PTM landscape across the human proteome.

Ultimately, we designed PTM-POSE for fast and reproducible analysis of experimental measurements of splice events/isoforms, allowing for generation of new, testable hypotheses. As a case study, we identified PTMs related to high ESRP1 expression in prostate cancer patients from The Cancer Genome Atlas (TCGA) using PTM-POSE, as this splicing factor correlates with poor patient outcomes. Through downstream functional analysis of differentially spliced PTMs, we found alterations to protein interaction and signaling networks that would not have been identified using exon-based tools alone, such as coordinated inclusion of SGK1 substrates. From this work, we have highlighted the importance of including post-translational modifications in the analysis of splicing and provided an easily implementable tool for annotating isoforms and splice events with PTMs and their functional consequences.

## Results

A fundamental challenge in studying the relationship between splicing and PTMs has been the lack of comprehensive PTM annotation across protein isoforms. Most PTMs in modern PTM compendia were discovered through mass spectrometry-based approaches that rely on trypsin-based proteolysis to yield MS-amenable peptide fragments. These measured peptides are then matched to specific proteins from a reference protein database, in a process dependent on the peptide matching algorithm and database used [26]. While isoform-aware databases have been developed to capture different protein isoforms [27], most PTM annotations from tryptic-fragment identification are assigned to the canonical UniProtKB isoform, unless the tryptic fragment can be uniquely assigned to an alternative isoform (Supplementary Figure 2). This limitation is reflected in current annotations - only 15.3% of alternative isoforms in UniProt have any recorded PTMs, compared to 93.1% of canonical isoforms, despite the fact that peptides containing modified residues could often be mapped to multiple possible isoforms.

To overcome this critical gap in knowledge, we integrated three key resources using a Python-based framework (Fig. 1). We combined genomic information from Ensembl (GRCh38.p14 version 111) providing experimentally-supported human protein-coding transcripts and exon locations [28], canonical and alternative protein isoform annotations from UniProt [29], and PTM data from ProteomeScout and PhosphoSitePlus [30, 31]. By integrating these resources, we created a comprehensive resource of PTMs mapped to their genomic location, including associated exons and transcripts. Given the genomic coordinates of a splice event, exon, and/or isoform, we can identify any PTMs mapped to that genomic region (Supplementary Figure 3). We refer to these as “prospective” PTMs, indicating that while the genomic location codes for a modifiable residue, there may not be direct evidence of the modification in the alternative isoform.

**Figure 1.**
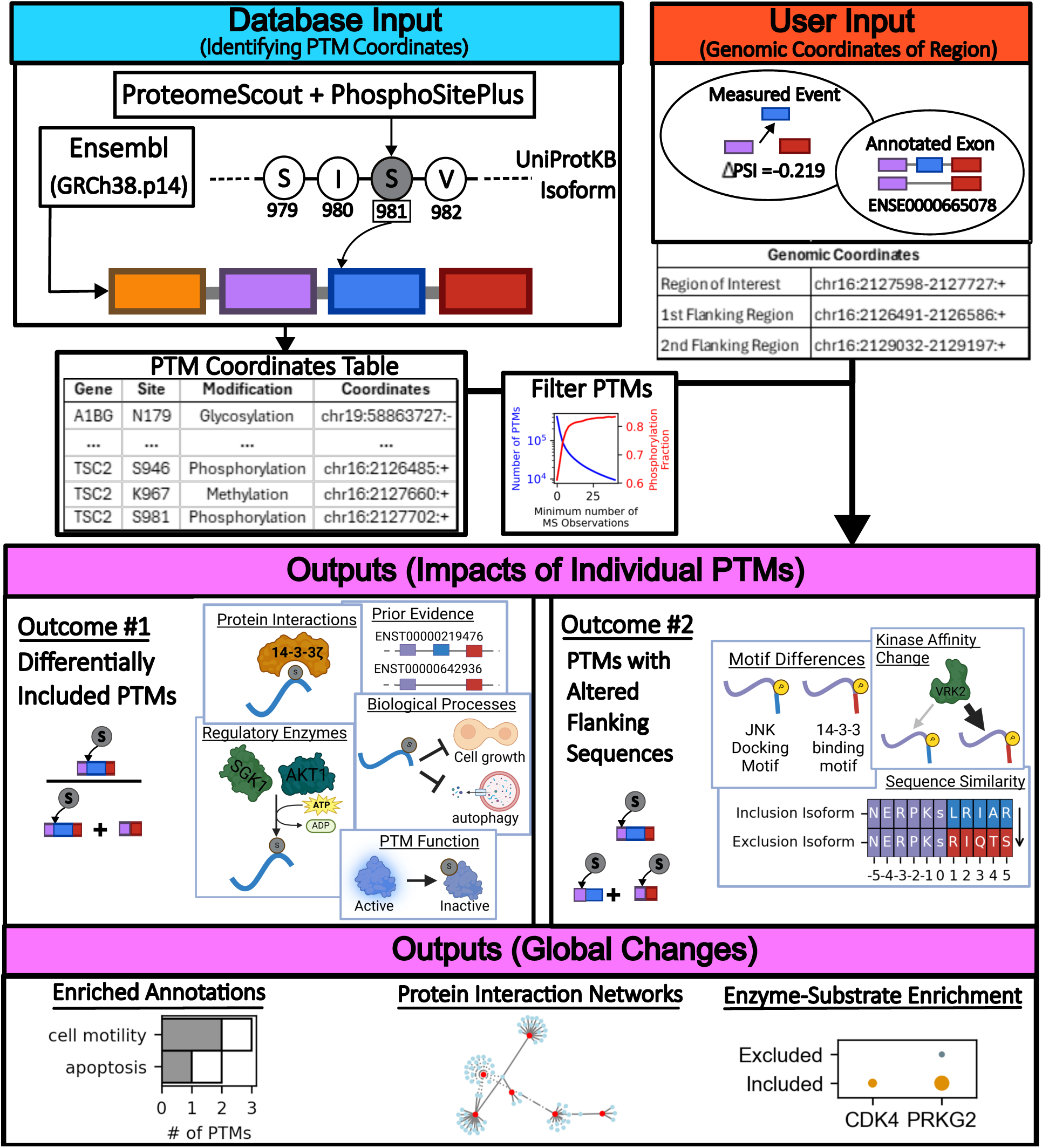
Mapping and Projecting Post-Translational Modification (PTMs) onto Isoforms and/or Splice Events with PTM-POSE. To expand the understanding of how PTMs are altered by alternative splicing, we first mapped all PTMs obtained from ProteomeScout [30] and PhosphoSitePlus [31] to their location in the corresponding exons, transcripts, and gene annotated in Ensembl, resulting in a comprehensive resource of PTM sites and their genomic locations. Then, given the genomic coordinates of a measured splice event or annotated transcripts as input, PTM-POSE identifies the PTMs associated with the genomic coordinates of the splice event/isoform. We defined two key mechanisms of PTM diversification by splicing based on the predominant events observed in our analysis: **(1)** the PTM is included in one isoform but not the other, or **(2)** the PTM is present in both isoforms, but the residues surrounding the PTM site were altered, which may influence the specificity of enzymes and binding partners. Once impacted PTM sites have been identified, PTM-POSE provides a suite of analysis modules focusing on three key areas of impact: i) changes to protein function or biological role, ii) changes to protein interactions, iii) or changes to interactions with regulatory enzymes like kinases.

To enable broad applicability, we developed PTM-POSE, an open-source tool that can project PTMs onto any measured splice event or exon, such as those quantified from RNA-sequencing data [32–34] (Fig. 1B). To facilitate use with a broad array of different data inputs, PTM-POSE only requires the genomic coordinates of a given event or exon in either hg19 or hg38 coordinate systems. These can be derived from quantified splice events from RNA-sequencing data, transcript annotations from databases like Ensembl, or any other data source that outputs genomic coordinates of splice events or exons. In addition, PTM-POSE provides the ability to filter prospective PTMs based on the degree of supporting evidence from ProteomeScout and PhosphoSitePlus, allowing users to focus on PTMs with higher confidence. However, for all analyses in this work, we opted to not filter any PTMs based on these metrics, as filtering tends to result in primarily only phosphorylation sites (Supplementary Figure 4).

Once splicing-related PTMs have been identified, PTM-POSE can be used to assess the functional consequences of these PTMs, integrating information from various PTM-focused databases and tools (see Supplementary Table 1 for full list). With these tools, users can assess the potential impact of PTM inclusion across isoforms or splice events in three main ways unique to PTM-POSE: i) known functions of individual PTMs based on annotations from databases like PhosphoSitePlus and PTMsigDB, ii) PTM-driven protein interactions, and iii) regulation of proteins by regulatory enzymes like kinases. Further, many analysis and plotting utilities are available in PTM-POSE for either close inspection of specific PTM sites or identification of globally over-represented PTM annotations and/or interactions. Finally, PTM-POSE allows users to directly compare to an exon-specific analysis tool, NEASE [15], to obtain a holistic view of potential splicing impacts. Importantly, we designed PTM-POSE for reusability and expansion as knowledge of both splicing and post-translational modifications grows.

In this work, using PTM-POSE, we performed systematic analyses to 1) characterize how alternative splicing impacts PTM inclusion and flanking sequences across the transcriptome, 2) examine modification-specific patterns of splicing control, and 3) demonstrate how splicing-mediated PTM regulation contributes to cancer-relevant protein regulatory networks. These applications highlight PTM-POSE’s utility in revealing new insights into how alternative splicing shapes protein function and regulatory networks through control of post-translational modifications.

### Projecting Prospective Post-translational Modification Sites onto Alternative Isoforms

Using PTM-POSE, we first performed a systematic analysis of PTM diversification across isoforms annotated in the Ensembl database, focusing on modifications that have at least 140 unique observations in the human proteome. We identified “prospective” PTMs associated with each annotated exon in Ensembl, allowing us to determine if alternative transcripts (not associated with canonical UniProtKB isoform) include some or all of the PTM-carrying exon. With this pipeline, we successfully expanded the annotations of PTMs on alternative isoforms across the proteome and transcriptome – 98.8% of prospective PTM sites identified in alternative isoforms from UniProt and/or Ensembl were not previously annotated in protein databases (Supplementary Figure 2C).

Given the successful projection of PTMs onto alternative isoforms, we next assessed the effect of splicing on PTMs based on a set of key possible effects (also described in Fig. 1A). First, a PTM may be included in an isoform, maintaining any potential interactions that have been shown for the canonical isoform. Second, a PTM could be excluded from an isoform altogether, removing any potential functional changes or interactions induced by the modification. Lastly, if a prospective PTM is included in the alternative isoform, it may contain a unique flanking sequence directly surrounding the PTM distinct from the canonical isoform, potentially leading to different interactions with regulatory enzymes or PTM-specific binding domains. The transcriptome-wide datasets, prospective PTMs, and regulation through splicing are freely available to the research community (see data availability section for details).

### PTMs are differentially included through alternative splicing

Based on the PTMs identified across Ensembl transcripts by PTM-POSE, we sought to evaluate how splicing impacts the availability of PTM sites in alternative isoforms and whether this relationship depends on the specific modification type. We began by examining the number of PTMs within Ensembl-defined constitutive exons – exons found in every transcript associated with a gene. Constitutive exons exhibited a lower density of PTM sites than non-constitutive exons, with the most densely populated exons tending to be non-constitutive (Supplementary Figure 5). This finding matches previous reports comparing tissue-specific exons to constitutive exons measured from RNA sequencing and provides further evidence that splicing plays a role in the inclusion of PTM sites in protein isoforms [5, 6].

To assess individual PTMs regulated by splicing, we defined “constitutive PTMs”, or PTMs that are included in all protein-coding transcripts of a gene and are therefore not controlled by splicing. Overall, 68.23% of PTM sites could be described as constitutive across Ensembl transcripts. Phosphorylation accounted for 64% (*n* = 248, 383) of all PTM sites assessed, with ubiquitination being the second most common at 24% (*n* = 93, 726) (Fig. 2A). Consequently, the overall constitutive rate likely reflects the rate of constitutive phosphorylation sites rather than PTMs in general, potentially skewing the representation of how different PTM types are diversified across isoforms. The constitutive rate for phosphorylation (68.2%) closely matches the overall rate, while other modifications, such as glycosylation (n = 16,920, rate = 74.6%), show greater deviations. When compared against null models where splicing is entirely random and independent of modifiable residues, we found that most modifications, except palmitoylation, exhibited slightly higher constitutive rates than expected by random chance, but the observed variability across modification types is unlikely to be observed by randomness alone (Fig. 2B). Constitutive rates ranged from 50.6% (Palmitoylation, 162 unique observations, *p ≤* 0.05) to 93.8% (Crotonylation, 192 unique observations, *p ≤* 0.05) (Fig. 2A,B, Supplementary Figure 6). Phosphorylation, sumoylation, hydroxylation and palmitoylation were among the modification types with the lowest constitutive rates. On the other hand, ubiquitination, glycosylation, crotonylation, carboxylation and sulfation had much higher constitutive rates than in the null models (Fig. 2B). PTMs found within structural domains were slightly more likely to be included in alternative isoforms, but this alone failed to explain the variation across modification types (Supplementary Figure 7). We also evaluated modification subtypes and found further variability in constitutive rates (Supplementary Figure 8). For example, glycosylation as whole exhibits a higher constitutive rate than most other modifications (74.6%). This is largely due to N-linked glycosylation (75.0%), while O-linked glycosylation has a constitutive rate closer to the average across all modifications (68.9%). Similarly, phosphoserines (67.6%) exhibited slightly lower constitutive rates than either phosphothreonine (68.8%) or phosphotyrosine (69.3%) sites. Together, these analyses reveal extensive control of PTM availability across protein isoforms while highlighting the importance of evaluating each modification type separately, even within the same modification class – as seen in the differences between O- vs. N-linked glycosylation and serine vs tyrosine phosphorylation.

**Figure 2.**
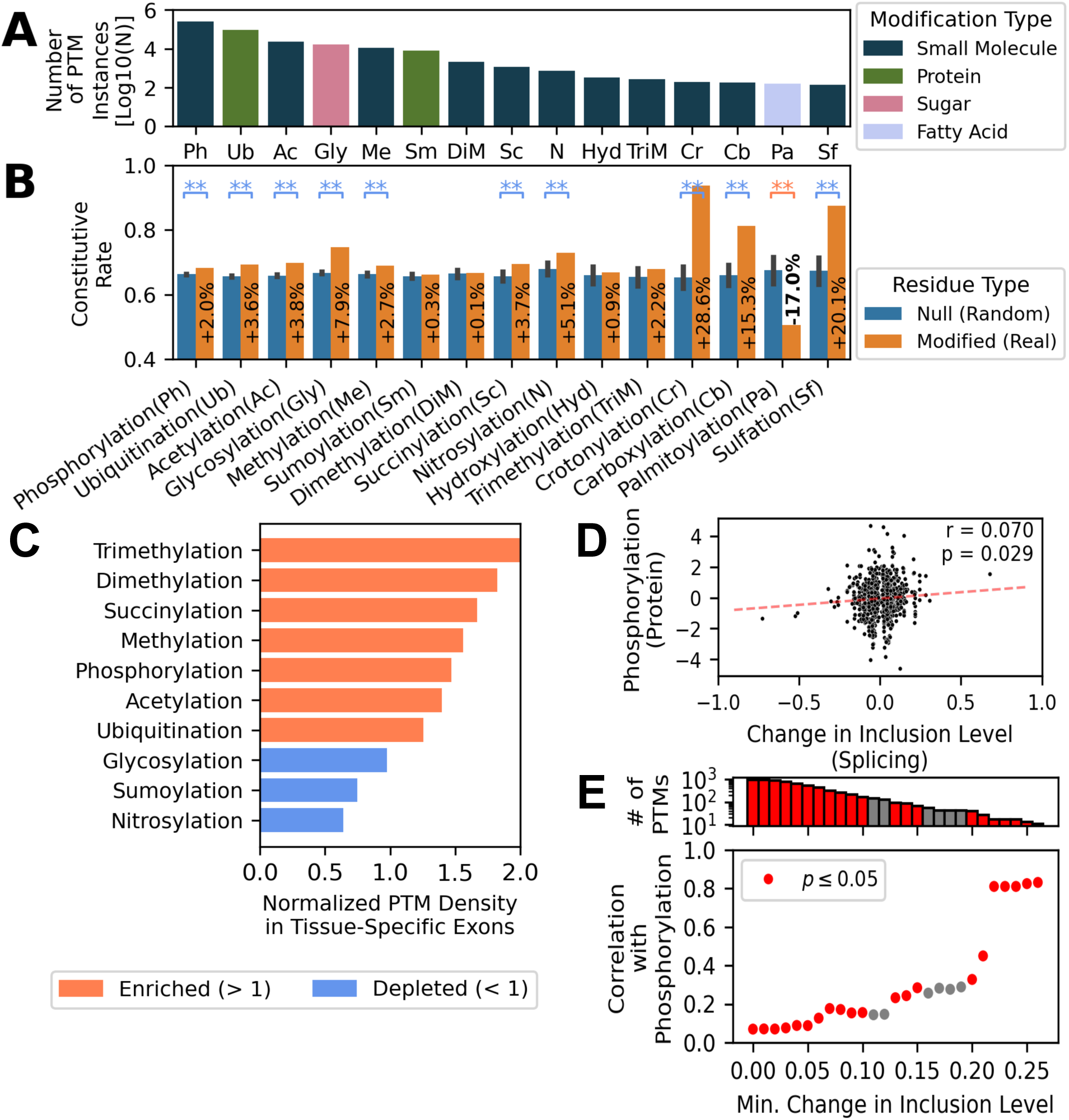
Splicing controls the presence of post-translational modifications in alternative isoforms. Modification-specific statistics of PTM inclusion across human alternative transcripts.**A)** The number of modifications documented in canonical UniProt proteins for each class of post-translational modification (min. 140 instances), colored by what type of molecule is being added. **B)** The fraction of modification sites found across all isoforms of a given gene, defined as the constitutive PTM rate. We compared rates to a null model generated by randomly selecting residues across the proteome and treating them as real PTM sites (see methods for details). Percent difference between null and real constitutive rates is indicated within the bars. **:*p ≤* 0.05. **C)** Density of each type of PTM in tissue-specific exons identified across three different publications [5] [35] [36], normalized by the density across the entire proteome. **D)** Comparison of the change in transcript inclusion for phosphorylation sites projected onto significant splice events (*FDR <* 0.05) quantified by rMATS [32] [33] and phosphorylation abundance measured by mass spectrometry. Data is based on measurements in lung cancer cells transduced with different KRAS variants (wild type, G12V mutation, and Q61H mutation). Each point represents a unique measured phosphorylation site. **E)** Correlation between transcript inclusion of phosphorylation sites and their measured abundance when filtering sites below a minimum threshold of inclusion.

Because we are harnessing data from all recorded instances of transcripts and PTMs from large databases, it is possible that the observed non-constitutive PTMs are due to either non-functional transcripts that do not ultimately produce proteins or PTMs that result from transcriptional and/or biological noise. To account for this possibility, we repeated the PTM-specific analysis with PTM-POSE using different definitions of functional transcripts, including functionality predictions from TRIFID [37]. As we implemented more restrictive functional criteria for transcripts, we observed a decrease in the number of isoforms per protein and a similar increase in the constitutive PTM rate, suggesting that the true constitutive rate is likely between 70-90% (Supplementary Figure 9A-D). Further, to ensure the observed constitutive rates are not driven by modification sites with limited evidence, we examined whether the constitutive rate changed when removing PTMs reported in fewer than 10 mass spectrometry experiments across literature and found that the constitutive rate actually decreased slightly from 68.1% (*n* = 246, 227) to 66.5% (*n* = 14, 352) (Supplementary Figure 9E). There also did not appear to be any relationship between constitutive rate and the number of different databases in which a site is recorded (Supplementary Figure 9F). Regardless of functional criteria used, the rates continued to retain similar variability across most modification types. Overall, these results support that there is extensive control of PTM availability across protein isoforms that is PTM-specific, although it is dependent upon what isoforms are ultimately expressed and functional within tissues.

To investigate the potential functional relevance of splicing-regulated PTMs, we identified enriched molecular functions and biological processes among PTMs controlled by splicing (non-constitutive) based on a Fisher’s exact test and annotations from PhosphoSitePlus [31]. Non-constitutive PTMs were found to be significantly associated with many functions important for regulating protein interactions and activity, including PTM sites important for intracellular localization (*n* = 2, 822, rate = 64.3%, *p* = 8*e −* 6) and regulating molecular associations (*n* = 5, 253, rate = 66.4%, *p* = 6*e −* 4), indicating that alternative splicing may rewire protein-interaction networks through exclusion of the PTM sites driving these protein interactions (Supplementary Figure 10, Supplementary Table 2). When looking at phosphorylation sites alone, these functions were also significantly enriched among non-constitutive sites (*p <* 0.05). In contrast, constitutive PTMs were only found to be enriched for sites regulating receptor desensitization (*n* = 139, rate = 83.4%, *p* = 6*e −* 4), and a larger than expected percentage of constitutive PTM sites had no known function (*n* = 379, 438, rate = 68.8%, *p* = 6*e −* 4). Similarly, constitutive PTM sites were not found to be enriched for any biological processes, while non-constitutive PTMs were enriched for a variety of processes including cell growth (*n* = 1, 778, rate = 65.6%, *p* = 0.016) and apoptosis (*n* = 1, 144, rate = 62.2%, *p* = 5*e −* 5) (Supplementary Figure 10). In addition, through PTM-POSE’s integration of kinase-substrate interaction data from PhosphoSitePlus [31] and RegPhos [38], we found that phosphorylation sites targeted by some kinases (RAF1, ERBB2, SRPK1) were more likely to be specific to particular isoforms, while substrates of other kinases (BUB1, GRK2, CAMK2A) tended to be present in all isoforms of their target proteins. (Supplementary Figure 10). Combined, this points to the idea that splicing control of PTMs across isoforms will impact protein function and the regulatory networks associated with the spliced protein.

Next, we examined if different types of splice events show distinct patterns in how they affect PTMs – such as skipped exons, alternative splice sites, or mutually exclusive exons. While the rate of splicing control of PTMs was modification-specific, most modifications were impacted by different splice events in similar proportions, with most excluded PTMs resulting from skipped exon events. Hydroxylation and sulfation sites were more frequently lost due to alternative splice sites in the alternative isoform compared to other modifications (Supplementary Figure 11A). In rare cases, mutually exclusive exon events may also impact the presence of PTMs, where an exon associated with an annotated PTM site is replaced with a mutually exclusive exon (MXE) with a similar sequence which may or may not still include the PTM. We identified 430 mutually exclusive exon events associated with PTM sites, using criteria defined previously by Pillman et al. [39]. These events encompass 153 different annotated PTM sites and 13 modification types (Supplementary Figure 11B,C). To determine whether the MXE replacing the canonical exon maintains the annotated PTM site associated with the canonical exon, we performed global sequence alignment of the mutually exclusive exon pairs and assessed whether the residue associated with the PTM and its surrounding sequence were present. We found that PTM sites were lost in 48.4% of cases, and the surrounding sequence was generally at least partially disrupted when the PTM was still present (Supplementary Figure 11D). This may indicate that mutually exclusive exon events are a mechanism by which PTM sites can be removed from alternative isoforms without significantly disrupting the overall structure of a protein.

While these results suggest an important role of splicing in controlling PTM inclusion, focusing on Ensembl-defined transcripts alone limits our understanding of the physiological relevance of these splicing-controlled PTMs, as it does not consider expression of transcripts within tissues. To assess differential inclusion of PTM sites across tissues, we used PTM-POSE to analyze a compiled set of tissue-specific exons identified across three different publications [5, 35, 36]. Overall, PTMs tended to be found at a higher density in tissue-specific exons than across the entire proteome, in line with prior results [5] [6]. As observed in our analysis of the entire transcriptome, PTM density in tissue-specific exons was largely dependent on the modification type. Glycosylation and nitrosylation exhibited lower density within tissue-specific exons than found across the entire proteome, matching their higher constitutive rate (Fig. 2C). Interestingly, sumoylation, which was observed to have low constitutive rates, was not found to be abundant within these tissue-specific exons. While the density of PTMs across tissue-specific exons indicates that the trends observed in the Ensembl transcriptome are unlikely to be solely an artifact of non-functional or rarely-expressed transcripts, these analyses still do not capture how alternative splicing of PTMs leads to changes at the protein level, where PTMs are relevant.

Given that phosphorylation is one of the more commonly spliced PTMs and high-throughput measurements of phosphorylation are more common than other PTMs, we focused on phosphorylation to assess how splicing affects modification presence at the protein level. In a prior study, expression of KRAS variants in human lung epithelial cells was shown to lead to changes in both phosphorylation patterns and alternative splicing events, providing an ideal case study for examining the relationship between splicing of phosphorylation sites and their measured abundance at the protein level [40]. Using RNA-sequencing analysis of the KRAS-expressing cell lines, we quantified splice events with rMATS [32, 33] and projected phosphorylation sites onto these splice events using PTM-POSE, yielding percent spliced in (PSI) measurements of PTM sites within transcripts for each experimental condition (wild-type, G12V, and Q61H KRAS variants). We then compared changes in inclusion of phosphorylation sites after KRAS variant expression to changes in phosphorylation abundance normalized for overall protein expression (Fig. 2D). In total, PTM-POSE identified an average of 4,243 phosphorylation sites with significant differences in inclusion after KRAS variant expression (*FDR <* 0.05) across the three experimental conditions. Of these, around 7.5% were also measured by mass spectrometry (*≈* 320 sites per condition), consisting mostly of phosphoserine sites (Supplementary Figure 12). Across all significantly spliced sites with corresponding phosphorylation measurements, we found a weak correlation between changes in inclusion level and changes in phosphorylation abundance (*r* = 0.07*, p* = 0.029*, n* = 966). However, as the difference in PSI increases, the correlation between the PSI difference and phosphorylation abundance also increases. In particular, a strong correlation emerges for sites with at least a 22% change in transcript inclusion, although this also reduces the available data (*r* = 0.67*, p <* 0.001*, n* = 23) (Fig. 2E). This suggests that while small changes in inclusion level may not lead to detectable changes in phosphorylation, differential inclusion of phosphorylation sites does lead to protein-level changes, particularly when focusing on PTMs with the largest changes in inclusion. For this reason, splice events and PTMs among the top inclusion changes should be prioritized to maximize the likelihood of measurable differences at the protein level (Δ*PSI >* 0.15 in this dataset, or the top 15% of changes). While the exact impact and scale of these changes on cell phenotype may still be unclear, the noticeable changes to protein modifications is likely to have downstream effects on protein function and interactions, highlighting the importance of considering PTMs associated with different isoforms and/or splice events.

### Alternative splicing affects PTM-regulatory sequences

PTMs are important components of signaling pathways, cellular communication, and protein interactions, all facilitated by direct interactions between PTMs and their corresponding modifying enzyme(s) and recognition domain(s). The specificity of these interactions is driven by the residues flanking the modification site, with the linear motif directly surrounding the PTM being a prerequisite for recognition of the PTM site [12, 13, 41]. There are several mechanisms by which alternative splice events could lead to changes in the flanking sequence of a PTM, including a skipped/retained exon adjacent to the PTM, an alternative splice site in a nearby exon or in the PTM-containing exon, or a mutually exclusive exon event (Fig. 3A). Using PTM-POSE, we investigated whether splicing may alter PTM function by changing its flanking sequence, defining the flanking sequence as the five amino acids on either side of the PTM, unless otherwise stated.

**Figure 3.**
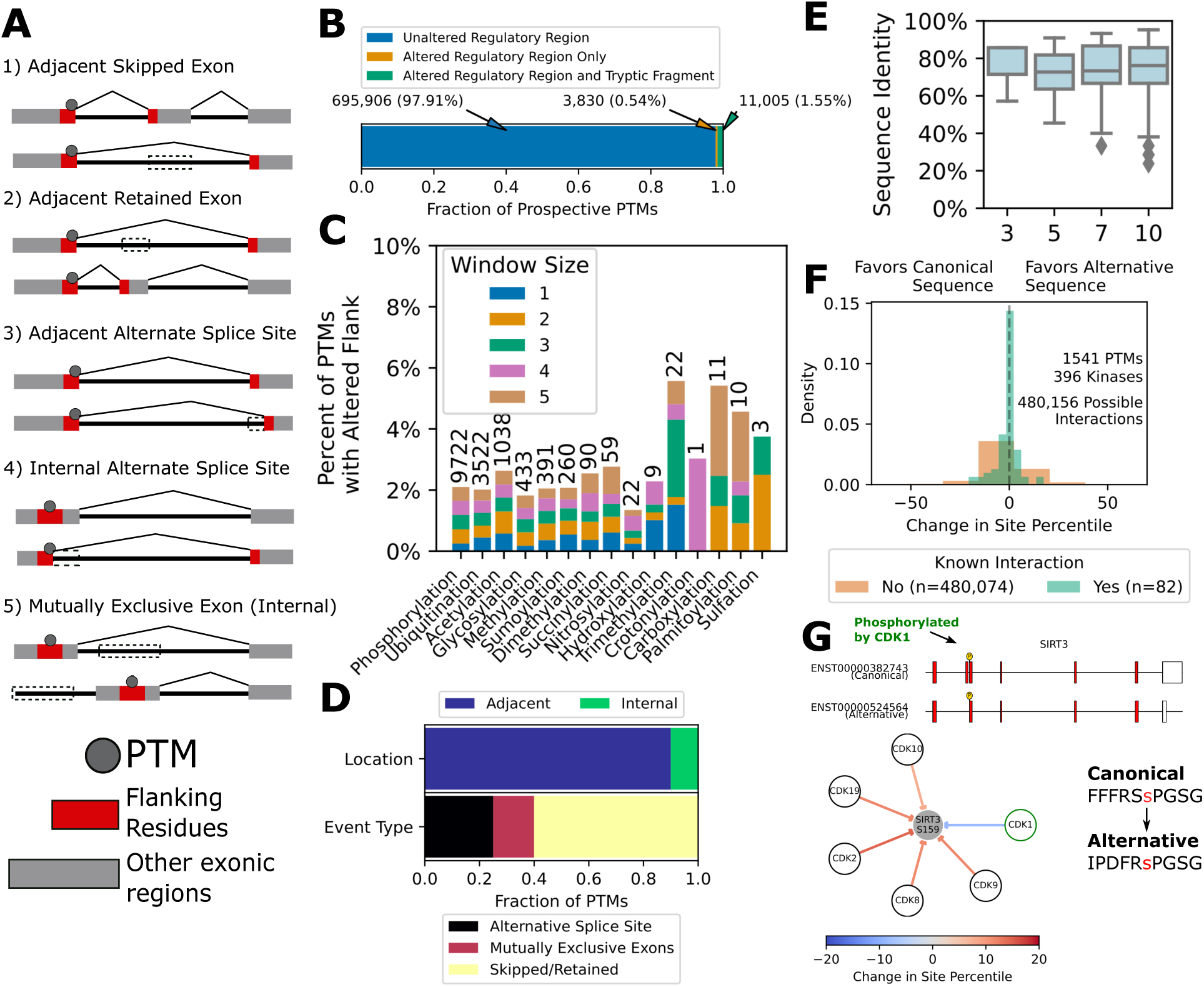
Alternative splicing rewires PTM-driven interaction networks by altering flanking sequences. **A)** Illustrative examples of the modes of alternative splicing that result in changes to the flanking sequence around a PTM, comparing two transcripts with differing flanking sequences (highlighted in red) surrounding the same PTM. **B)** Number of prospective PTMs found in alternative isoforms with an unchanged flanking sequence (+/- 5 amino acids), an altered flanking sequence but unchanged tryptic fragment, or both an altered flanking sequence and tryptic fragment. **C)** Fraction of prospective PTMs with altered flanking sequences, broken down by modification type, with the total number of events indicated above each bar. The proximity of the change to the PTM is denoted by the different bar colors, ranging from immediately next to (1) or up to 5 amino acids away from the PTM on either side. **D)** Breakdown of which events described in panel A most commonly lead to altered regulatory regions. **E)** Sequence identity between differing flanking sequences in the canonical and alternative isoforms, using different window sizes of amino acids on either side of the PTM. **F)** Change in the likelihood of kinase interaction from the canonical to the alternative flanking sequence, based on predicted site percentiles derived from Kinase Library software [41]. We only considered PTM sites with an altered flanking sequence and conserved tryptic fragment (orange bar in panel B) that also had at least one known kinase interaction annotated in PhosphoSitePlus [31] **G)** An example of an altered flanking sequence surrounding the phosphorylation site at S159 in SIRT3. The change in Kinase Library site percentile between the canonical and alternative isoform of SIRT3 for various cyclin-dependent kinases are shown, with positive changes (red arrows) indicating a preference for the alternative isoform, while negative changes (blue arrows) indicating a preference for the canonical isoform.

When comparing flanking sequences of a PTM site in canonical and alternative isoforms, we found that 2.09% of prospective PTM sites had changes to their flanking residues (Supplementary Table 3). Importantly, 25.8% of these sites have the same tryptic fragment as the canonical isoform, indicating that 0.54% of prospective PTM sites have altered regulatory regions that would not be detected by trypsin-mediated MS (Fig. 3B). Across the possible altered regulatory regions, acetylation, trimethylation, carboxylation, and palmitoylation exhibited the highest rate of alteration (Fig. 3C). When compared against the null model, we found that the high alteration rate could not be explained by randomness alone (Supplementary Figure 13). In the majority of cases, altered flanking sequences are caused by changes to the adjacent exon, rather than the PTM-containing exon itself, usually via a skipped/retained exon (Fig. 3D). Both glycosylation and nitrosylation, which also had higher constitutive rates, showed lower rates of altered flanking sequences which could be attributed to the distribution of amino acids across the transcriptome (Supplementary Figure 13).

We further explored how splicing affects specific residue positions in the flanking sequences and the potential consequences of these changes. Due to the nature of splicing-induced changes, most regulatory regions are altered predominantly on one side of the modification’s flanking sequence, and the farther from the PTM the more likely changes are to occur(Supplementary Figure 13). Among the *≈* 2% prospective PTMs with altered linear motifs, 7.5% of these PTMs (2,381 prospective PTMs) have a change in the +/-1 residue immediately adjacent to the PTM site, which has the potential to significantly impact enzyme or binding domain interactions (Fig 3C, Supplementary Figure 13). For example, proline-directed kinases are a large subset of serine/threonine kinases with a strong preference for a proline in the +1 position, which includes cyclin dependent kinases (CDKs) and ERK [41]. Similarly, the CDY family of proteins contain a chromodomain that binds to methylated lysines with a strong preference for lysines that immediately follow an arginine [42]. In both cases, the remaining residues in the regulatory window serve to drive the specificity and strength of the interaction, with more minor effects. When comparing the sequence identity between flanking sequences in the canonical and alternative isoform, we found that regardless of the regulatory window size chosen (3, 5, 7, or 10 amino acids on either side of the PTM), flanking sequences generally still maintained *>* 60% sequence identity, suggesting that splicing-induced regulatory sequence changes preserve most information in the motif (Fig. 3E). Altogether, these results indicate that splicing can alter regulatory sequences in ways that could impact PTM-specificity and recognition, but suggest that it is unlikely to fully disrupt the modifiability of a PTM.

We next sought to understand how changes in the regulatory sequences might affect PTM regulation and protein interactions. Given the limited data regarding isoform-specific binding or enzymatic activity, we used the kinase motif enrichment tool Kinase Library to compare the predicted affinity of various serine/threonine kinases for both canonical sequences and alternative regulatory sequences identified by PTM-POSE [41]. We focused on phosphorylation sites with an altered flanking sequence and a conserved tryptic fragment, as these are the modifications most likely to be missed by trypsin-mediated MS and be misannotated on the canonical isoform. In total, we analyzed 328 phosphotyrosine events (286 unique sites) and 1,484 phosphoserine/threonine events (1,255 unique sites), 82 of which had a known annotated kinase interaction. Interestingly, in many cases, flanking sequence changes had diverse impacts on both annotated kinase interactions (kinases that had been previously observed to phosphorylate a given site) and on kinases not annotated as a regulatory kinase: decreased likelihood in some cases, increased in others, and in an equal number of cases had no effect on the predicted likelihood (Fig. 3F). Kinases not known to interact with a particular phosphorylation site exhibited more extreme cases than the annotated, known kinase interactions. In some cases, the predicted ranking for unannotated kinases differed by more than 100 ranks between the canonical and alternative isoform, while predicted changes in the annotated interaction tended to be more subtle. However, there were also cases where the annotated kinase’s predicted interaction differs substantially in the alternative isoform. For example, SIRT3, a deacetylase that regulates energy metabolism, contains an activating phosphorylation site at S159 that has been shown to be phosphorylated by CDK1. In an alternative isoform of SIRT3, a skipped exon event results in a unique regulatory sequence preceding S159 (Fig. 3G). While the proline in the +1 position remains present (key feature of proline-directed kinases like CDKs), the surrounding changes result in a decreased likelihood of phosphorylation by CDK1 while increasing affinity for other CDK family members like CDK2, suggesting a potential difference in the regulation of phosphorylation at S159, moving away from CDK1 in the alternative isoform (Fig. 3G). Although limited by the scarcity of known enzyme interactions, the analyses enabled by PTM-POSE revealed how splicing-induced changes in PTM flanking sequences can diversify a protein’s regulatory networks by altering enzyme or binding domain specificity.

### Profiling ESRP1-related splicing in prostate cancer with PTM-POSE

We wished to apply PTM-POSE to examine how splicing-control of PTMs alters protein function and regulatory networks in disease. Aberrant splicing is involved in several diseases, including cystic fibrosis, muscular dystrophy, and cancer [47–51]. Our global analysis found that non-constitutive PTM sites were enriched for sites important to carcinogenesis, suggesting that changes to splicing patterns may influence the presence of functionally important PTM sites for the development and progression of cancer (Supplementary Figure 10). Further, many splicing factors regulating pre-mRNA splicing have been implicated in altering cancer progression, such as epithelial splicing regulatory proteins (ESRPs), which are responsible for splicing epithelial-specific isoforms [52]. In prostate cancer, ESRP1 is commonly amplified (8% of patients from the TCGA cohort) and has been reported to lead to worsened prognosis when overexpressed [51, 53, 54]. The majority of studies focus on ESRP1’s role in the epithelial to mesenchymal transition (EMT), with low expression of ESRP1 leading to a more mesenchymal and invasive state, although there have been reports that this does not occur in some tissues [52, 55–58]. Previous literature has also identified a strong link between increased ESRP1 expression, high copy numbers, and chromosomal deletions in prostate cancer, although the mechanism for this remains unclear [51, 53]. Hence, ESRP1 expression in prostate cancer represents an interesting application of integrated PTM and splicing analysis with PTM-POSE. We hypothesized that ESRP1-mediated splicing may lead to changes in PTM inclusion and flanking sequences that drive changes in protein function and interactions that contribute to prostate cancer progression.

To identify whether ESRP1 expression in prostate cancer results in differential inclusion or altered flanking sequences of specific PTM sites, we used PTM-POSE to analyze splice events measured in ESRP1-low (*z ≤ −*1, *n* = 61) and ESRP1-high (*z ≥* 1, *n* = 68) prostate cancer patient samples from The Cancer Genome Atlas (TCGA), identified based on normalized mRNA expression. By applying PTM-POSE to splice events quantified in TCGASpliceSeq [34], we could compare both exon and PTM utilization in transcripts across samples within the prostate cancer cohort, defined by the percent spliced in (PSI) rates of each exon and PTM. PTM-POSE identified a total of 1,115 PTM sites that were significantly differentially spliced based on ESRP1 expression, representing 0.5% of PTMs found in the dataset (Mann Whitney U test: *p ≤* 0.05, *r ≥* 0.3). Of these, 468 PTM sites were excluded and 647 PTM sites included when ESRP1 is highly expressed (Fig. 4A, Supplementary Table 4). Phosphorylation makes up the the majority of differentially included modifications, but we also identified several acetylation, ubiquitination, and methylation sites affected by ESRP1 expression (Supplementary Figure 14). Across the 6.8% of genes in the dataset associated with significant splice events as a result of ESRP1 expression (557 genes), 54% of these genes contained PTMs impacted by ESRP1-associated splicing, with around half of impacted exons containing PTMs (Fig 4B). A subset of the identified PTMs (107) were also associated with splice events occurring after ESRP1 knockdown in H358 lung cancer cells, where PTMs included in ESRP1-high patients were decreased after knockdown. This includes phosphorylation sites found in TSC2, CTNND1, CD46, and MAP3K7, indicating that we successfully captured at least some ESRP1-mediated splicing in these patient samples [60](Supplementary Figure 14). Additionally, PTM-POSE identified 334 PTMs, or 0.2% of PTMs assessed, that had altered flanking sequences due to differential inclusion of adjacent exonic regions as a result of ESRP1 expression (Mann Whitney U *p ≤* 0.05, *r ≥* 0.3)(Fig. 4B). In sum, PTM-POSE identified 1,489 different PTM sites across 562 different exons impacted by differences in ESRP1 expression, indicating extensive regulation of PTMs by ESRP1.

**Figure 4.**
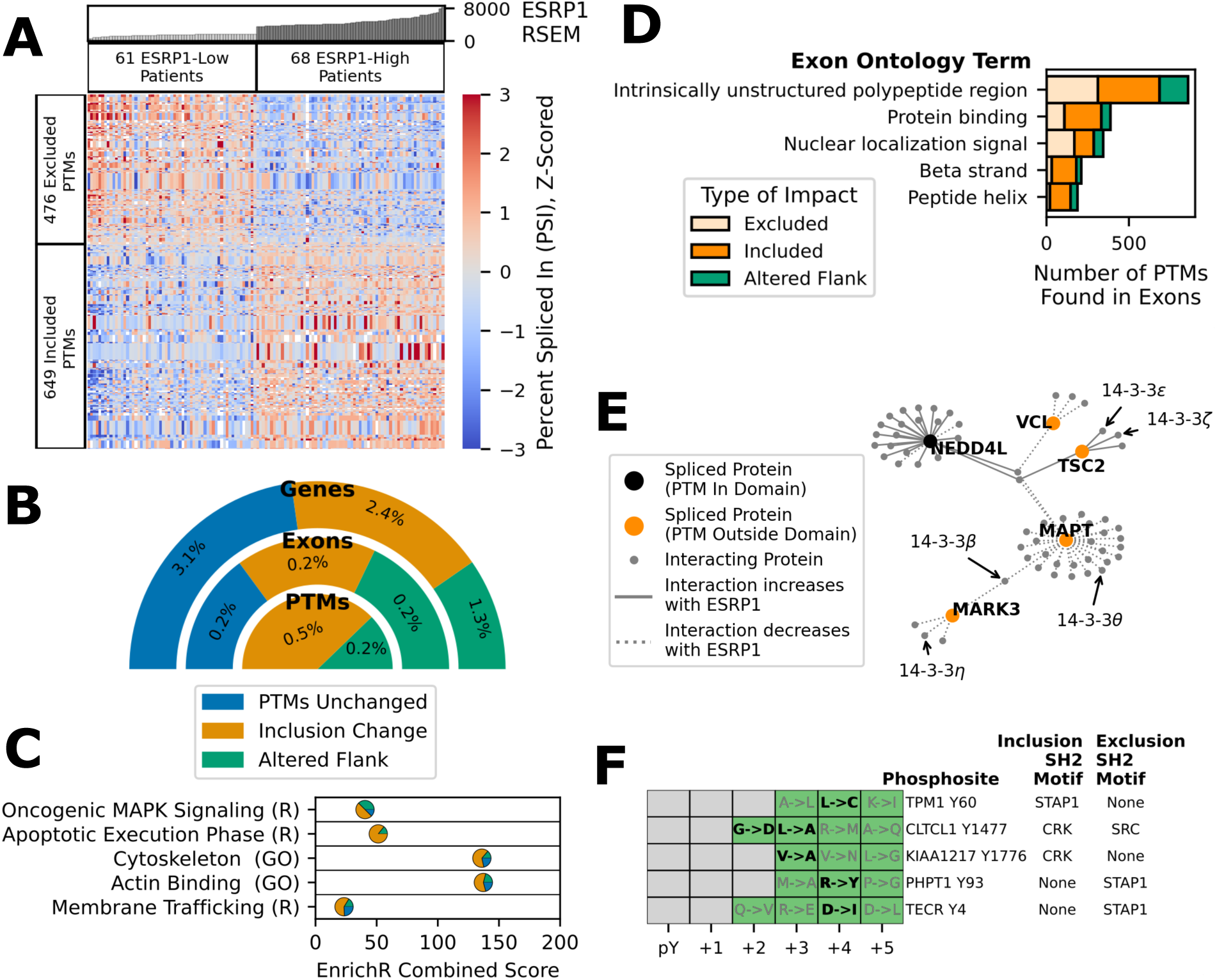
ESRP1 drives changes to protein function and interactions in prostate cancer by altering the presence and flanking sequences of specific PTM sites. Using data from The Cancer Genome Atlas and SpliceSeq, we explored the relationship between ESRP1 expression and splicing-controlled PTM sites in prostate cancer [34] [59]. **A)** Patient-specific percent spliced in (PSI) of individual PTM sites that were found to be differentially included between ESRP1-low patients (*z ≤ −*1) and ESRP1-high patients (*z ≥* 1). **B)** Breakdown of the fraction of genes, exons, and PTMs impacted by splicing, including whether the impact is due to changes in inclusion or alterations to the surrounding flanking sequence of a PTM. **C)** Selected gene sets from Gene Ontology (GO) [43] and Reactome (R) [44] that are enriched among spliced genes with PTMs that are impacted (orange and green groups from Panel B). Each point indicates the fraction of spliced genes associated with the gene set belonging to each group in Panel B. **D)** Exon Ontology terms [14] that are associated with the highest number of impacted PTMs, broken down by the type of impact (excluded, included, or altered by ESRP1) **E)** Subset of the PTM-associated interactions impacted by differential inclusion of PTMs due to ESRP1 expression, all of which were impacted by inclusion differences greater than 16%. Interactions are based on data from PhosphoSitePlus [31], RegPhos [38], and PTMcode [45]. Orange nodes indicate proteins with spliced PTMs outside domains, and black nodes indicate proteins with spliced PTMs within domains. **F**) Phosphorylated tyrosine sites with different flanking sequences based on ESRP1 expression, which only match the preferred linear motif of select SH2 domains depending on whether an adjacent exon is included in the protein isoform. Each row indicates a different phosphorlated tyrosine site and the specific residue change is denoted as ”Inclusion AA -*≤* Exclusion AA”. Changes that specifically impact the motif are bolded and in black.

### Identifying PTM-associated processes with PTM-POSE

To identify the primary roles of PTMs impacted by ESRP1-mediated splicing, we first looked at the the general functions of genes associated with spliced PTMs using EnrichR [61, 62]. From this analysis, we found that genes associated with spliced PTMs were enriched for cytoskeletal and actin-binding proteins, with over 75% of the spliced cytoskeletal genes containing PTMs impacted by ESRP1-related splicing, leading to a higher EnrichR score than when considering all spliced genes (Fig. 4C, Supplementary Figure 15). This matches expected results given ESRP1’s role in regulating cell motility and the epithelial-to-mesenchymal transition. Intriguingly, many pathways were only significantly enriched when considering genes with impacted PTMs rather than all spliced genes, including a subset of genes that were also involved in the apoptotic execution phase pathway (CASP8, ADD1, MAPT, PTK2, SPTAN1) (Fig. 4C, Supplementary Figure 15).

Beyond gene-level functional information, PTM-POSE allows for deeper exploration of the function of individual PTM sites using annotations from PhosphoSitePlus [31]. Spliced PTMs related to cell motility were among the most common processes associated with regulated PTMs (6 identified), but they did not indicate a consistent change in motility (some changes would inhibit motility, others would promote), suggesting that either ESRP1 does not drive motility changes in prostate cancer or that these changes are not PTM-driven (Supplementary Figure 16). We also observed several individual PTM sites that are associated with cell growth, cell cycle regulation, and apoptosis (Supplementary Figure 16). Several of these PTMs, but not all, suggest increased proliferation and decreased apoptosis including CRK Y221 (induces apoptosis, decreased inclusion with ESRP1 expression) and PAK4 S291 (induces cell growth, increased inclusion with ESRP1 expression). However, there were also other identified PTMs, such as EEF1A1 S21 (induces apoptosis, increased inclusion in ESRP1 expressing cells), highlighting the importance of context on PTM function and the need to consider all altered proteins and PTMs holistically.

### PTM-POSE reveals altered PTM-driven interactions by ESRP1-mediated splicing

When looking across the transcriptome, non-constitutive PTM sites were significantly associated with regulating molecular associations and protein interactions (Supplementary Figure 10). Matching this finding, among ESRP1-related PTMs identified by PTM-POSE, the most common functions were related to regulating molecular associations and intracellular localization (24 and 7 PTMs, respectively) (Supplementary Figure 16). Further, PTM-containing exons impacted by ESRP1-mediated splicing were in regions of protein binding and nuclear localization signals, suggesting that ESRP1 control of PTMs may lead to altered protein interaction and regulatory networks (Fig. 4D). The majority of these PTMs were found in exonic regions associated with intrinsically unstructured regions of the protein, rather than distinct domains (59.1%) (Fig. 4D). Given the importance of the PTMs in facilitating protein interactions, we used PTM-POSE to integrate PTM-associated interactions from PhosphoSitePlus [31], RegPhos [38], and PTMCode [45]. A total of 476 PTM-associated interactions were impacted by ESRP1-mediated splicing out of 25,510 potential interactions, with many PTMs facilitating these interactions falling outside of specific structural domains (Fig. 4E, Supplementary Figure 17). Notably, when comparing these findings to interactions that could be identified using exon-level information alone through NEASE (domains, linear motifs, etc.) [15], only 67 of the 476 PTM-driven interactions were also found by NEASE, highlighting the benefit of analyzing impacted PTM sites in conjuction with other information (Supplementary Figure 17). Upon ESRP1 expression, we noted several proteins overrepresented in the PTM-associated interaction network (hypergeometric test: *p ≤* 0.05), including decreased interactions of ABL/SRC interacting proteins (CRK, ABI1), increased interactions of several ribosomal proteins (RPL6, RPS14, RPS3A), and variable changes to interactions of cytoskeletal proteins (MAPT, CTNND1). The increase in PTM-associated interactions of ribosomal proteins aligns with increased proliferation often observed among ESRP1-expressing cancer cells [52]. Several ribosomal proteins, including RPL6 and RPS14, have also been linked to chromosomal instability and DNA damage response, although it is not clear how the affected phosphorylation, methylation and ubiquitination sites may regulate these processes [63, 64]. When prioritizing interactions impacted by inclusion changes greater than 16% (85th percentile of PTM-associated changes) to maximize possibility of splicing impacting availability of these PTM sites, we identified a subnetwork of splicing-regulated proteins consisting of NEDD4L, TSC2, MARK3, and MAPT. Among these proteins, we noted several impacted interactions with 14-3-3 proteins, important scaffolding proteins involved in regulation of many cell signaling pathways that are commonly dysregulated in cancer [65] (Fig. 4E). Combined with the aforementioned apoptosis and cell growth-related PTM sites, these findings suggests that ESRP1-mediated splicing of PTMs may drive worsened prognosis in prostate cancer by rewiring protein interactions that regulate growth-related signaling pathways and protein translation, leading to increased cell growth and chromosomal instability, rather than through the more commonly studied EMT pathways.

In addition to differential inclusion of PTM sites, we also used PTM-POSE to assess cases where changes to flanking sequences may alter protein interactions by disrupting preferred linear motifs of specific binding domains. We found similar patterns of disruption as observed in our transcriptome-wide analysis, where most changes to the flanking sequence retained 60-80% sequence identity after exclusion of the adjacent exon (Supplementary Figure 18A-C). Among the altered flanking sequence events, we found many events that disrupted potential targets of various binding domains with motifs annotated in the Eukaryotic Linear Motif (ELM) database, including the motifs for 14-3-3 proteins (bind phosphorylated serines and threonines) and three different SH2 domains (bind to phosphorylated tyrosines) (Supplementary Figure 18D-E) [66]. For example, the SH2 domain of CRK preferentially binds to proline, leucine, isoleucine, or valine in the +3 position, which are changed to an alanine for CLTCL1 Y1477 and KIAA1217 Y1776 when ESRP1 is expressed (Fig. 4F). PTMs with disrupted motifs for 14-3-3 proteins showed more varied patterns of disruption, but most impacted genes were related to cytoskeletal genes like MARK2 and SORBS2, which have been shown to interact with 14-3-3 protein epsilon based on interaction data from STRING (Supplementary Figure 18) [67]. Combined with the disrupted interactions by altered inclusion of PTMs, these results highlight how splicing control of PTMs by ESRP1 can rewire whether and how proteins interact with each other, such as the interactions of 14-3-3 proteins.

### ESRP1 expression is correlated with SGK1 signaling in prostate cancer

While some functional differences between isoforms can be gleaned without PTM-specific information from PTM-POSE (based on structural domains, linear motifs, etc.), the presence of PTMs also helps define how the protein isoform is regulated by modifying enzymes. Kinases, the enzymes responsible for phosphorylation, are often implicated in cancer progression and are the targets of many cancer therapies [70]. Using PTM-POSE, we can identify substrates of specific kinases that are differentially included as a result of ESRP1 expression, indicating potential coordinated changes to how these kinases behave in these cellular systems. Among the known kinase-substrate interactions from PhosphoSitePlus [31] and RegPhos [38], we found an increase of multiple substrates of AKT1, SGK1, and PRKACA (median Δ*PSI* = 10.8%), a decrease of ABL1 substrates (median Δ*PSI* = *−*5.4%), and several substrates of MAPK1/3 that were altered due to ESRP1 expression (median Δ*PSI* = *−*10.3%) (Fig. 5A, Supplementary Figure 19). However, given that kinase-substrate information tends to be sparse (less than 5% of phosphorylation sites have an annotated kinase), we employed PTM-POSE’s integrated kinase-substrate enrichment analysis adapted from a previously developed kinase activity algorithm, called KSTAR [46], to determine if there were significant inclusion changes to kinases’ substrates. We found that substrates of SGK1 and other SGKs were enriched among phosphorylation sites with increased inclusion in ESRP1-high patients from TCGA cohort, while AKT1 substrates were actually more commonly found in ESRP1-low patients.

**Figure 5.**
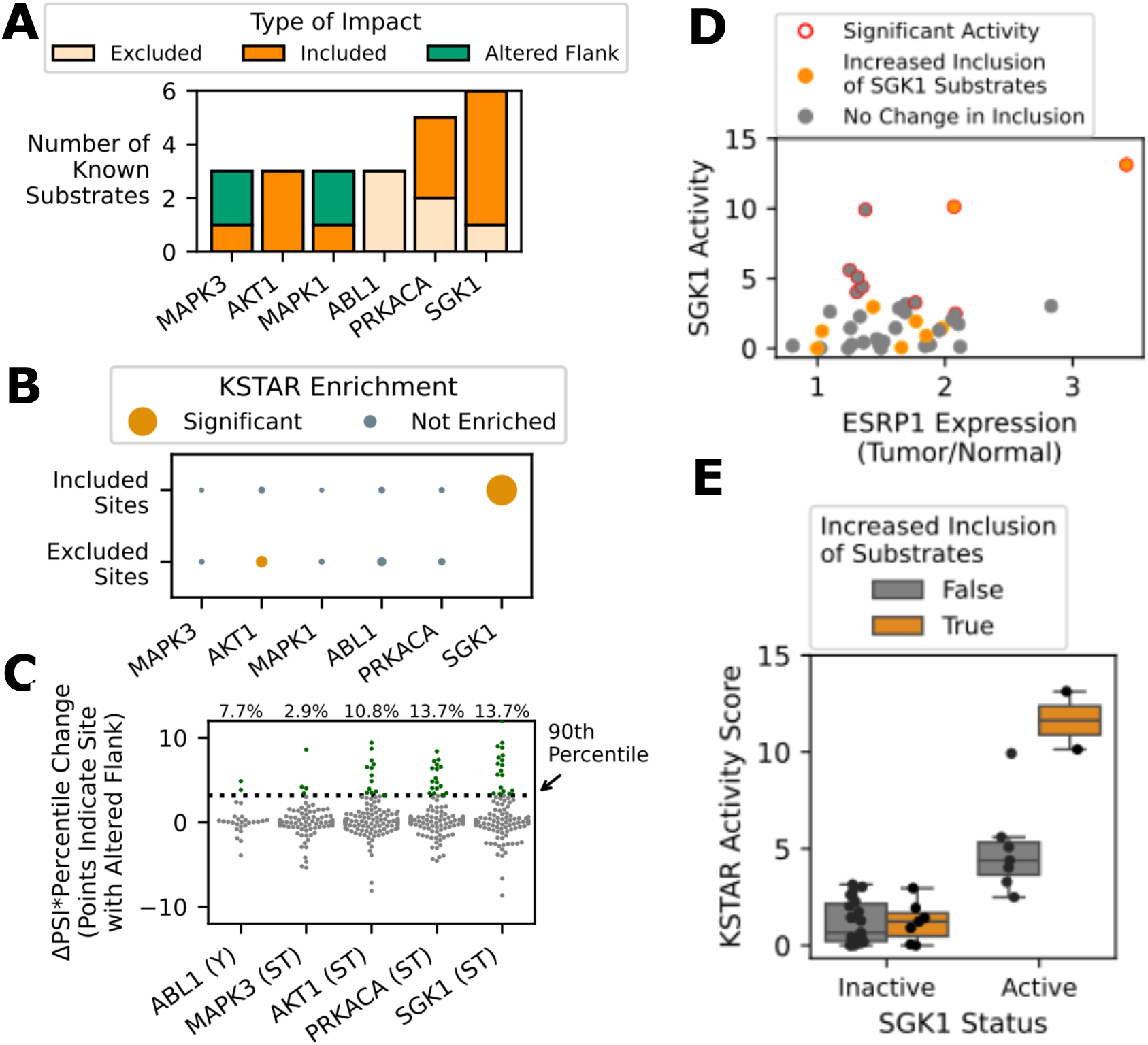
ESRP1 rewires kinase signaling networks through changes to their downstream targets. Given the phosphorylation sites identified to be impacted by ESRP1 splicing in prostate cancer in Figure 4, we assessed whether these sites tended to be targets of specific kinases. **A)** Kinases with the highest number of differentially included substrates, based on either known kinase-substrate interactions from PhosphoSitePlus [31] and RegPhos [45] **B)** Enrichment of kinase-substrate interactions among phosphorylation sites that are either included or excluded when ESRP1 is highly expressed. Kinase-substrate enrichment was performed using an adapted version of kinase activity inference algorithm, called KSTAR [46]. The size of the dots represent the strength of enrichment, with color indicating significance. **C)** Change in the likelihood of a kinase-substrate interaction due to changed flanking sequences. Interaction change is defined as the change in kinase library percentile between the inclusion and exclusion isoforms multiplied by the change in percent spliced in (PSI), such that a positive change indicates the kinase is more likely to interact in ESRP1-high patients and a negative change indicates that is less likely to interact. We have highlighted interations that fall in the 90th percentile in green (most likely to increase upon ESRP1 expression). **D)** Comparison of predicted SGK1 activity from phosphoproteomic measurements by KSTAR [46], ESRP1 expression, and enrichment of SGK1 substrates in sites with change in inclusion greater than 10% in individual CPGEA patients [68, 69]. **E)** Difference in predicted SGK1 activity scores as a function of SGK1 substrate inclusion, whether looking at patients with inactive or active SGK1.

Given that kinase specificity is partially determined by the linear motifs surrounding a phosphorylation site, we also assessed if altered flanking sequences around phosphorylation sites may change the likelihood of kinase interactions. Based on the change in PSI and predicted scores from Kinase Library, SGK1 and PRKACA exhibited the highest variability in the likelihood of interacting with phosphorylation sites containing altered flanking sequences as a result of ESRP1 expression, whereas MAPK3 actually seemed to be least impacted by flanking sequence changes (Fig 5C). However, it should be noted that many of the same phosphorylation sites observed increased preference for multiple of the assessed kinases, with only DOCK7 S958 and SVIL S1177 being uniquely increased in preference for SGK1. Altogether, the targets of SGK1/2/3 appear to be most impacted by ESRP1-mediated splicing in prostate cancer patients.

### Validating the impact of splicing on phosphorylation and kinase activity in prostate cancer

Although we found that certain kinases’ substrates were more commonly included in ESRP1 expressing prostate tumors due to splicing differences, the actual phosphorylation of these targets depends on protein expression of the translated product and the activity of a kinase, among other factors. Despite the relationship between SGK1 substrate splicing and ESRP1, it was not clear how these changes ultimately influence SGK1 substrate phosphorylation, activity, and signaling in ESRP1-expressing tumors. Unfortunately, the TCGA prostate cohort lacks proteomic measurements by mass spectrometry and the phosphorylation sites measured by Reverse Phase Protein Array (RPPA) were not found to be impacted by splicing. Therefore, to evaluate the relationship between spliced phosphorylation sites, measured phosphorylation, and kinase activity readouts, we turned to a smaller prostate cancer cohort from the Chinese Prostate Cancer Genome and EpiGenome Atlas (CPGEA) cohort. This dataset contains both RNA sequencing and phosphoproteomic measurements of normal and tumor tissues from 40 patients [68, 69]. We applied PTM-POSE to splice events quantified by rMATS [33] in CPGEA samples and found 1,038 PTMs correlated with ESRP1 expression, including 267 PTMs that were also found to be related to ESRP1 expression in the TCGA cohort (24% of TCGA sites) (Supplementary Figure 14). As with the TCGA dataset, ESRP1-correlated PTMs were enriched for SGK1 substrates, suggesting similar trends in the CPGEA cohort and the CPGEA dataset can be used to validate the initial findings in the TCGA cohort (Supplementary Figure 19).

The CPGEA dataset’s matched proteomic data also provided an opportunity to validate whether splicing changes identified by PTM-POSE correspond to protein-level changes in PTM abundance. When comparing the spliced phosphorylation sites to their measured abundances by mass spectrometry in each patient, 1.9% of the sites with sufficient variability in transcript inclusion across patients in the CPGEA cohort (*PSI range ≥* 0.4, *PSI standard deviation ≥* 0.05) were measured by mass spectrometry, with 58 of the ESRP1-correlated PTMs being measured (Supplementary Figure 20). We found that inclusion changes exhibited weak correlations with phosphorylation abundance (*r* = 0.02, *p <* 0.05), similar to what was observed in the KRAS overexpression study discussed in Figure 2. The largest changes in inclusion were more likely to correspond to the proteomic measurements (Δ*PSI ≥* 0.4, maximum correlation = 0.68, Supplementary Figure 20). This suggests that even in noisier contexts of patient data, the splicing-related changes in PTM inclusion can still be detected at the protein level, providing a good dataset to assess the impacts of splicing on kinase signaling.

Kinase activity is commonly inferred from phosphoproteomic measurements of a kinase’s substrates, with active kinases exhibiting a larger number of their substrates being phosphorylated in a sample [46]. As a result, kinase activity predictions are readouts of both substrate availability and enzymatic activity. To assess how splicing of a kinase’s substrates relates to changes in kinase signaling, we predicted kinase activity in prostate tumors relative to matched healthy samples from CPGEA using KSTAR [46, 68, 69]. When comparing the proteomic-based kinase activity measurements to a similar enrichment analysis of differentially spliced phosphorylation sites from each patient, kinases with differentially included substrates were more commonly found to have significant activity (29% of kinases were active when substrates were enriched among included sites, 20% otherwise)(Supplementary Figure 21). In addition, active kinases with increased inclusion of their substrates tended to exhibit higher activity scores (*p <* 0.05), supporting the idea that splicing of phosphorylation sites does not lead to an active kinase but can increase a kinase’s degree of activity in a patient tumor by increasing substrate availability (Supplementary Figure 21).

When assessing the impact of ESRP1 on kinase activity, three kinases had significant correlations with ESRP1 expression across the 40 CPGEA patients: SGK1, PIM1, and ABL2 (*p <* 0.05,Supplementary Figure 21). Given that SGK1 substrates were also identified as influenced by ESRP1-mediated splicing, we dug deeper into the relationship between ESRP1 expression, SGK1 expression, SGK1 substrate splicing, and SGK1 activity (Fig. 5D). In total, there were twelve patients that exhibited increased inclusion of SGK1 substrates by splicing (*p <* 0.05, increase in availability), two of which were also predicted to have elevated levels of activity (*FPR ≤* 0.05, increase in both substrate availability and phosphorylation). Notably, these two patients also had high ESRP1 expression and the highest activity scores of all patients with predicted SGK1 activity (Fig. 5E). To determine whether SGK1 activity might be related to ESRP1-driven changes in expression rather than splicing of its substrates, we examined SGK1 transcript expression (not measured in proteomic data). Interestingly, SGK1 transcript expression is inversely correlated with ESRP1 protein expression and does not significantly correlate with predicted SGK1 activity (Supplementary Figure 21). Altogether, this suggests that ESRP1 may facilitate increased SGK1 signaling through splicing of its substrates and an increase in substrate availability, with the requirement that SGK1 is already active in the patient tumor. Importantly, without a PTM-specific analysis of splicing changes with PTM-POSE, it would not have been possible to identify the potential link between ESRP1 and SGK1, highlighting the utility of PTM-POSE in identifying novel hypotheses between splicing and regulatory networks.

## Discussion

As highlighted in a recent review, even small changes (loss of linear motifs, PTMs, allosteric effects, etc.) between protein isoforms can lead to functional differences and phenotypic changes in cells [71]. While various methods and resources have been developed to assess potential isoform-specific functions and interactions, consideration of PTMs and their function has been limited (only report presence of PTM) or absent from these resources [14–22]. In this work, we developed a pipeline to map PTMs to their genomic location and project them onto alternative isoforms and splice events, culminating in a publicly available tool we call PTM-POSE. Using PTM-POSE, we first performed a comprehensive analysis of how splicing affects PTMs and their linear motifs across the Ensembl transcriptome, revealing several key principles. First, we showed that inclusion/exclusion of PTMs by splicing is extensive. Second, we demonstrated that regulatory or protein interaction networks can be diversified through both differential inclusion and alterations in the linear motifs of PTMs. Lastly, we established that different types of PTMs should be analyzed separately, given their distinct biochemical properties, regulation, measurement methods, and relationships to splicing. The abundance of documented phosphorylation sites compared to other modifications means that analyses grouping all PTMs together mainly reflect splicing’s impact on phosphorylation. However, in our analysis, each modification type exhibited specific patterns in constitutive rates, tissue specificity, flanking sequence regulation, and even the types of splicing events that alter the PTMs.

When applying PTM-POSE to specific experimental contexts, we also found measurable relationships between differential splicing of phosphorylation sites and their abundance at the protein level in two separate contexts (KRAS expression, prostate tumors), despite abundance being dependent on numerous other factors including protein translation, enzyme expression/activity, and protein degradation. Across both datasets, the top *≈* 15% of inclusion changes across PTM sites showed the strongest agreement with measured phosphorylation, although this corresponded to different PSI thresholds (Δ*PSI ≥* 0.15 in KRAS expression dataset, Δ*PSI ≥* 0.4 in CPGEA dataset). These findings establish PTM-POSE’s ability to identify biologically relevant splicing changes that affect PTM sites, while also highlighting the importance of considering context-specific thresholds that can be used to prioritize hypotheses surrounding differentially included PTMs. Importantly, though, we view these findings as a preliminary exploration of the relationship between splicing and PTMs, with PTM-POSE allowing for continued study in other experimental contexts in which splice event or isoform information is available.

Our global and experiment-specific studies highlighted a major limitation stemming from the lack of PTM annotation, including context-specific functional roles and regulatory connections of individual PTM sites. Less than 5% of PTMs are annotated with additional functional context in PhosphoSitePlus [31], with phosphorylation being one of the better annotated PTM types. This inherently limits our current understanding of the full scope of the impact of isoform-specific PTMs on protein functional diversity, as well as why certain types of PTMs are more likely to be regulated by splicing. Even still, PTM-POSE enabled discovery of larger patterns between splicing and PTM-driven regulation that would not have been identified with other splicing analysis tools. For example, we found that substrate-based predictions of kinase activity were related to coordinated splicing of the kinase’s substrates in prostate cancer patients, with active kinases tending to have larger activity scores when their substrates were differentially included by splicing. In particular, PTM-POSE revealed a previously unknown connection between SGK1 activity and ESRP1-mediated splicing in prostate cancer that merits further investigation, a finding that was enabled through PTM-specific analyses of splicing.

Although we have lowered the barrier for integrating PTMs into splicing anaylsis by developing tools for annotating experimentally observed splice events with PTMs and their functions, experimental validation at the protein level remains challenging. The vast majority of differentially spliced PTMs occur in ways that make them effectively indistinguishable amongst isoforms by traditional mass spectrometry workflows. For the same reason that PTM databases predominantly annotate canonical isoforms, current experimental approaches cannot attribute peptide-wise modifications to specific protein isoforms. Even in cases where both isoform information (such as from RNA-sequencing) and PTM abundance data are available, we found limited overlap between the two data types, constraining our ability to make broader conclusions. To overcome some of these limitations, future work could explore using custom splice-aware databases for peptide matching, incorporating PTM-containing peptide sequences identified from transcriptomic data [27]. In addition, complementary analysis with other splicing tools that assess splice events effects on isoform expression and function, such as those that predict nonsense mediated RNA decay (NMD), could further improve identification of functionally relevant isoform-specific PTMs and the overlap between RNA-sequencing and phosphoproteomics. Alternatively, information from PTM-POSE could be used to design splicing-specific panels of phosphorylation sites for measurement by targeted phosphoproteomic approaches, similar to ones developed to analyze specific signaling pathways or cancers [72, 73]. For example, to assess how ESRP1 expression influences SGK1 activity in prostate cancer cells, a targeted approach could specifically measure the SGK1 substrates impacted by splicing in ESRP1 expressing prostate cancer that were not measured by untargeted proteomics of CPGEA samples, such as TSC2 S981 and NEDD4L S448.

On the other hand, this work also highlights a novel perspective on interpreting PTM measurements – that the relative quantification of PTMs by mass spectrometry could reflect of changes in overall modifiability of target peptides across isoforms. For example, increases in measured phosphorylation, often attributed to an increased phosphorylation by a kinase on a single protein isoform, might instead reflect increased exon inclusion of that modifiable residue, rather than an overall increase in kinase activity. This insight suggests that careful examination of splicing patterns using PTM-POSE could help interpret changes in PTM abundance detected by proteomic experiments.

Together, this study suggests key avenues for advancement including improving annotations of modifications (their function and interaction partners) and further developing the experimental and computational approaches to connect modifications to specific isoforms. Additionally, we recognize that the field is rapidly improving and expanding in identification of new modification sites and transcript isoforms. Given this continual growth, we have developed and provided the computational pipeline developed in this work as an easily implementable tool for annotating RNA-sequencing datasets with PTM sites, as well as enabling new global analyses as our knowledge base expands.

## Methods

### Data acquisition and processing

To identify genomic coordinates of PTMs across the transcriptome, exon and coding sequences for protein-coding genes belonging to GRCh38.p14 build (version 111) of Ensembl were downloaded using the Biomart web interface. Additional exon, transcript, and gene meta information was downloaded from Biomart using the pybiomart package. Transcript sequences were derived from exon sequences and exon rank information, and the coding sequence was matched to its location in the transcript. Transcripts producing a protein isoform with fewer than twenty amino acids were removed from analysis. Post-translational modifications and other protein-level information associated with UniProt proteins were downloaded from ProteomeScout [30] and PhosphoSitePlus [31]. Protein sequences from Ensembl, ProteomeScout, and PhosphoSitePlus were compared and any discrepancies were removed from analysis. Transcripts that coded for the same protein sequence were collapsed into a single isoform entry. TRIFID functional scores associated with each transcript were downloaded from the APPRIS database [74].

RNA-sequencing fastq files for the KRAS variant overexpression study from Lo et al. were downloaded from the Sequence Read Archive (SRA) using the SRA toolkit and the following reads from the following accessions: SRR11247605, SRR11247606, SRR11247607, SRR11247608, SRR11247609, SRR11247610, SRR11247599, SRR11247600, SRR11247601, SRR11247602, SRR11247603, SRR11247604 [40].

For ESRP1-related splicing analysis, the SpliceSeq splicegraph and percent spliced in (PSI) values for individual events measured in at least 75% of patients were downloaded for the prostate cancer cohort from the TCGASpliceSeq database (https://bioinformatics.mdanderson.org/TCGASpliceSeq/) [34]. All other data corresponding to the prostate cancer cohort was downloaded directly from the cBioPortal (mRNA expression, clincal data, etc.) [59]. For the prostate cancer cohort from the CPGEA project, the proteomic, phosphoproteomic, and associated meta data was downloaded from the National Omics Data Encyclopedia (NODE) at the accession number OEP004998 [69]. RNA-sequencing fastq files for the forty patients from CPGEA with matched proteomic and phosphoproteomic data were downloaded from the Genome Sequence Archive for Human (GSA-human) at the project accession number PRJCA001124.

### Mapping PTMs to the Genome and Projecting them onto Alternative Isoforms in Ensembl

In order to map post-translational modifications to their location in the genome, coding sequences from Ensembl were first translated into the corresponding amino acid sequences using Biopython. Coding sequences that did not start with a start codon or had incomplete codons were removed from analysis. Based on the position of the PTM site in the protein and the location of the coding sequence in the transcript, the transcript location of the codon responsible for the modifiable residue was obtained and the corresponding exon containing that codon identified. Using the genomic coordinates of the exon of interest, the PTM could then be mapped to its location in the genome in the hg38 coordinate system, creating a repository of PTMs and their genomic coordinates. The pyliftover package was then used to convert from hg38 coordinates to both hg19 and hg18 coordinate systems. We have compiled the pipeline for mapping PTMs onto the genome in a python package called ExonPTMapper (https://github.com/NaegleLab/ExonPTMapper), and all data associated with this work are freely available (see data availability section).

To assess PTMs across isoforms annotated in Ensembl or measured from RNA-sequencing data using PTM-POSE, we identified mapped PTMs that are located within the genomic coordinates of an individual exon or splice event. To ensure that the PTM was correctly projected and was not disrupted by a frame shift, we verify that the codon associated with the projected PTM site codes for the correct residue. When projecting PTMs onto alternative transcripts in Ensembl (Figures 2 and 3), we also considered the rare case where a PTM site exists at the splice junction and is coded for by two exons. In these cases, we checked for conservation of one of the two contributing exons in the alternative transcript, then validated that the residue remained unchanged. We call the PTMs that were successfully projected on an alternative transcript or splice event a “prospective” PTM site, as the majority of these sites have not been experimentally validated. See Supplementary Figure 3 for a toy example of this process. Using this pipeline, we have developed a open-source python package, called PTM-POSE, for projecting PTMs onto any splice event or isoform with available genomic coordinates.

### Identifying ”Splice Events” relative to the Canonical Isoform

When thinking about the different types of splice events that may impact PTMs, we focused predominantly on skipped exon events, alternative splice sites, and mutually exclusive exon events, as these are the events that can lead to loss of PTM sites and are a direct result of splicing. Given that most PTM-level information used in this work is documented on canonical UniProt isoforms, splice events were defined relative to the transcripts associated with the canonical UniProt isoform. Genes for which there was not a clear canonical Uniprot isoform were excluded from this analysis. For each exon associated with the canonical isoform, we first checked to see if the Ensembl exon ID matched any exons in the alternative transcripts. In cases where there were no matching IDs, we compared the genomic location of the exon in the canonical isoform to the genomic locations of exons in the alternative transcript. From this, we identified cases where 1) an alternative exon had the same genomic coordinates (“Conserved”), 2) no alternative exons have any overlapping genomic coordinates (“Skipped”), 3) or an alternative exon has partially overlapping genomic coordinates (“Alternative Splice Site”). For “Conserved” cases, we checked to see if the protein sequences coded for by the exon within both the canonical and alternative isoform were the same, or if they had been altered due to a frameshift or alternative promoter. These cases were removed from analysis, so that focus was specifically on splicing mechanisms. For exons that were identified as “Skipped” in an alternative transcript, we assessed whether there was potential for a mutually exclusive exon event, using criteria defined by Pillman et al. [39]. In brief, we identified exons in the alternative transcript as candidate mutually exclusive exons if they were adjacent to the skipped exon location and not found in the canonical isoform. We then defined true mutually exclusive exons as those that had a length difference of no more than 20 amino acids (60 nucleotides) and a sequence similarity of 15% or more. Sequence similarity was determined through global pairwise alignment between the canonical exon and MXE candidate, using gap penalties of -10 when introducing gaps and -2 for extending gaps. Alignment was performed using Biopython [75].

### Calculating PTM and flanking sequence conservation across isoforms

For each known PTM site, we identified all alternative protein-coding transcripts associated with a gene and calculated the fraction of transcripts found to contain the PTM site. PTM sites that were found in all isoforms of a given gene were defined as “constitutive”. Given that multiple transcripts can code for the same protein sequences, we collapsed transcripts with matching protein sequences into a single isoform entry prior to calculating conservation. We could then calculate a “constitutive rate” for different modification types, which is the fraction of PTMs of that modification type that could be defined as constitutive. For cases in which transcripts were filtered out based on different functional criteria, potentially dysfunctional transcripts were removed from analysis prior to calculating constitutive rates (Supplementary Figure 9).

In order to compare flanking sequences between canonical and alternative isoforms in the Ensembl genome, we obtained the flanking sequences based on the projected location of the PTM in the alternative isoform, usually considering the five amino acids on either side of the PTM unless mentioned otherwise. To obtain tryptic fragments, we identified the nearest lysine and/or arginine residues to the PTM that are not proceeded by a proline, as this is the preferential cut sites for the enzyme trypsin [76]. We then compared the flanking sequences between the canonical and alternative isoform and identified cases in which the flanking sequence did not match (at least one amino acid was different). When calculating the overall rate of alteration for these flanking sequences, we only considered PTM sites that were successfully projected onto the alternative isoform of interest (i.e. it does not incorporate PTMs that are excluded from the isoform entirely).

In order to identify cases of changed flanking sequences from splice events (such as those identified in KRAS variant study and prostate cancer samples), we projected PTMs onto the flanking exons (exons immediately upstream or downstream of the spliced regions) of each significant splice event. For PTMs that were within fifteen base pairs of the splice junction (five amino acids), we used the Ensembl REST API to extract the DNA sequence associated with both the flanking regions and the spliced region. We then stitched together two DNA sequences, one which includes the spliced region (inclusion region) and one which excludes the spliced region (exclusion region). We translated these DNA sequences into amino acid sequences, using the PTM location to define the reading frame for translation (assume no frame shifts introduced). If extracted flanking sequences were too short (less than five amino acids on either side), the PTM was removed. We then compared the flanking sequences of the inclusion and exclusion regions and identified cases in which the flanking sequence was different.

### Generating the null splicing model for comparison with patterns of modifications

In order to determine if the observed splicing control of PTMs was unique to modifiable residues and not a general property of all residues, we developed a null model in which “modifiable” residues are randomly distributed across the transcriptome. For each modification type, we randomly sampled an identical number of residues as there are the modification of interest, with the same distribution of the specific amino acids that can be modified. For example, for the phosphorylation null model, we randomly sampled a total of 223,659 residues (131,236 serines, 54,358 threonines, 38,025 tyrosines), the same number of phosphorylation sites that have been observed in the proteome. With the random sample of residues, we then repeated the same analysis as done for the true modifiable residues (constitutive rate, flanking sequence identity, etc.). We repeated this process 100 times for each modification type to create a null model distribution of constitutive rates and altered flanking sequence rates. Finally, we considered the rate statistically significant if the observed value was found to have a higher or lower value than all but five or fewer of the random null models (*p ≤* 0.05).

### Identifying tissue-specific PTM sites

Tissue-specificity data was downloaded from three different publications which used different approaches to identifying for tissue-specific exons and/or transcripts [36] [5] [35]. In cases in which only tissue-specific transcripts were reported, we compared the two transcript isoforms to identify tissue-specific exons, excluding alternative splice site events. Across all tissue-specific exons, density of PTMs, defined as the number of PTM sites per 1000 residues, were calculated by counting the number of PTM sites whose genomic coordinates fall within the tissue-specific exons and dividing that number by the amino acid length of each exon. Amino acid length excluded residues which were found at splice junctions and were coded for by two exons. PTM density was then normalized to the density of PTMs found across the entire proteome, defined as the 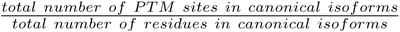. Thus, a normalized density greater than one indicates a higher density of PTM sites then expected relative to the entire proteome.

### Quantifying splice events from RNA-sequencing data with rMATS-turbo

To identify relevant splice events from RNA-sequencing data, when not provided by the original study, we utilized rMATS-turbo (v4.1.1) [69]. Briefly, we downloaded the relevant fastq files for each sample and aligned reads to the human reference genome (NCBI GRCh38 build) using STAR (v.2.7.9a) [77]. Alignment was performed using parameters recommended in the rMATS-turbo publication. The aligned reads were then used as input to rMATS-turbo using default parameters, except for specifying the expected read length and allowing for variable read length of the RNA-seq reads.

### Predicting changes in kinase interactions using Kinase Library

To assess how changes to a PTM’s flanking sequence may impact which proteins interact with that site, we turned to a kinase motif enrichment tool, Kinase Library [41]. To get scores from Kinase Library, we first restricted our analysis to prospective PTMs in alternative isoforms with altered flanking sequences (based on five amino acids) and a matching tryptic fragment. In total, we scored a total of 1812 different events. For each, we used the Kinase Library web interface to score the flanking sequence in both the canonical and alternative isoforms, and then extracted the ’site percentile’, which indicates where the phosphorylation site ranks relative to all other phosphorylation sites in the phosphoproteome scored for that kinase. We then calculated the change in site percentile from the canonical to alternative sequence for the entire transcriptome, or between the inclusion and exclusion flanking sequences for specific splice events in prostate cancer. For the latter, we further normalized the change in site percentile by the change in percent spliced in for the given exon to capture the effect of the splicing event on the kinase interaction.

### Identifying PTM sites related to ESRP1 expression in prostate cancer

In order to identify PTM sites specific to individual prostate cancer patients from The Cancer Genome Atlas (TCGA), “percent spliced in” values (PSI) specific to each patient were downloaded from the TCGASpliceSeq database, which indicate the fraction of transcripts within a given patient for which the exon of interest is included [34]. PSI values were calculated using SpliceSeq, an algorithm designed to predict alternative splicing patterns from RNA sequencing data, like the ones generated by TCGA [34]. While the majority of this work relies on Ensembl hg38 coordinates, TCGASpliceSeq reports exons in terms of hg19 coordinates, so genomic coordinates of PTM sites were converted into the hg19 coordinates system using the pyliftover python package. From these genomic locations, we projected PTMs onto SpliceSeq exons using the streamlined version of the procedure used for all Ensembl transcripts, PTM-POSE, which only requires the genomic coordinates of the splice event quantified by SpliceSeq. Next, we downloaded patient-specific expression of ESRP1 mRNA across the from cBioPortal [59]. We also downloaded survival data and alteration data for ESRP1 and PTEN from cBioPortal. ESRP1-high and low patients were defined as patients with ESRP1 mRNA expression greater or less than one standard deviation from the mean expression across the TCGA cohort, respectively. Finally, we compared inclusion of SpliceSeq exons across the ESRP1-low and -high groups and identified statistically significant differences using a Mann Whitney U test and Benjamini-Hochberg FDR correction (*p ≤* 0.01, *r ≥* 0.25). PTM sites projected onto these differentially included exons were then also defined as differentially included. To identify PTMs that have altered flanking sequences as a result of each splicing event, we instead identified the PTMs associated with flanking exon regions (regions immediately adjacent to spliced region). For each PTM found within 5 amino acids (or 15 nucleotides) from the splice junction, we translated the flanking sequence based on whether the spliced region was included or excluded, assuming no frame shifts had been introduced into the protein isoform, which we found to be a rare event across the entire transcriptome.

Due to lower sample numbers from the CPGEA prostate cancer cohort, instead of binning patients into low and high ESRP1 expression groups as done for TCGA, we performed correlation analysis to identify ESRP1-related events, treating both tumor and matched healthy samples as unique. First, we filtered splice events quantified by rMATS to include only events measured in at least 20 samples and some variability across samples (*range ≥* 0.4, *standard deviation ≥* 0.05). After filtering, we calculated the Spearman correlation between ESRP1 protein expression and PSI values for each splice event. ESRP1-related events were then identified as those with significant correlation after Bonferonni correction (*r ≥* 0.3, *p ≤* 0.05). PTM sites associated with these events were then defined as ESRP1-related PTMs.

### PTM- and gene-level enrichment analysis

PTM-site specific enrichment analysis was performed utilizing annotations downloaded from PhosphoSitePlus [31] and Exon Ontology [14]. Enrichment for a function or process was assessed using a Fisher’s Exact test, where the background population either included all PTM sites in canonical isoforms (as in Fig. 2) or all PTMs identified in the TCGA prostate cancer SpliceSeq dataset (as in Fig4). P-values were corrected using Benjamini-Hochberg false positive correction [78]. Gene-specific enrichment analysis was performed using the Enrichr wrapper from the gseapy python package [79] [61] [62]. Gene sets from Gene Ontology and Reactome were used [43] [44]. The background population was defined as all genes identified in the TCGA prostate cancer SpliceSeq dataset (as in Fig. 4).

To identify kinases with enriched substrates regulated by ESRP1 (Supplementary Figure 19), we utilized kinase-substrate networks of known connections from PhosphoSitePlus [31] and RegPhos [38] or predicted connections from KSTAR [46]. In both cases, a one-tailed hypergeometric test was used to assess statistically significant enrichment of a kinase’s substrates across ESRP1-regulated PTMs, with all phosphorylation sites identified in the TCGA SpliceSeq data used as the background population. For the KSTAR networks, the median p-value enrichment was extracted across the ensemble of 50 predicted kinase-substrate networks to obtain a final enrichment value.

### Constructing a PTM-associated interaction network

In order to identify protein-interactions potentially impacted by splicing-controlled PTMs, we pulled interaction information from PhosphoSitePlus [31], RegPhos [38], and PTMcode [45]. For each resource, we converted the information into common UniProt accession numbers and then combined the three resources into a single interaction network, with each node being a specific PTM site facilitating the interaction. We then identified the differentially spliced PTMs from the TCGA prostate cancer cohort that could be identified in the interaction network. From this network, we condensed the PTM sites into single protein nodes and defined as either a gained interaction (PTM sites are included in ESRP1-high patients) or lost interaction (PTM sites are excluded in ESRP1-high patients). For the few cases where the PTM was indicated to disrupt the interaction, we defined the interaction as anti-correlated with inclusion. The proteins with the highest number of edges (interactions gained or lost) were then reported.

To determine the impacts of altered flanking sequences on protein interactions, we utilized linear motif classes annotated in the Eukaryotic Linear Motif database (ELM), including the preferred linear motifs for common ligand binding domains [66]. ELM reports each classes motif as a regular expression, which we then used to search for the presence of each motif within the five amino acid flanking sequence of a PTM site, repeating this for the flanking sequence that occurs during the inclusion and exclusion events. We identified cases in which a specific linear motif was found in only one event (i.e. either when the adjacent exon is excluded or when it is included, but not both). Further, for linear motifs that specifically bind to a type of PTM, such as SH2 domains or 14-3-3 proteins, we checked to make sure the motif aligned with the position of the PTM site in the flanking sequence, as ELM motifs do not account for modified residues.

### Kinase activity prediction using KSTAR

Kinase activity of prostate cancer patients from the CPGEA was done using the KSTAR python package [46]. For each patient, we extracted the ratio of tumor to matched healthy tissue for each PTM site. For serine/threonine kinase activity predictions, we used a threshold of 1.5, 150 random experiments, and 100 significance trials. This led to a median of 833.5 phosphorylation sites used as evidence of kinase activity for each patient. For tyrosine kinase activity predictions, due to the limited number of measured tyrosine sites in the dataset, we used all sites measured in a patient as evidence, 150 random experiments, and 100 significance trials. This led to a median of 34.5 tyrosine sites used as evidence for prediction. After running KSTAR, mann whitney activities and false positive rates were extracted and plotted for each patient.

## Data and Code Availability

All data generated for this work has been deposited in Figshare at https://doi.org/10.6084/m9.figshare.24969993. The computational pipeline used to map PTMs onto transcripts and their genomic location and project them onto alternative transcripts is freely available at the following github repository: https://github.com/NaegleLab/ExonPTMapper, v0.2.0. Further, we have developed a streamlined version this code for use with most splicing quantification tools, called PTM-POSE https://github.com/NaegleLab/PTM-POSE, v0.1.0. All additional code and analysis done for this work can be found at a separate github repository here: https://github.com/NaegleLab/PTM_Splicing_Analysis, v3.0.0.

## Acknowledgements

We would like to thank Noah Perry and Mete Civelek for their guidance and input throughout the development of this work. Additionally, we would like to thank Ben Jordan for early mentorship of undergraduates working on this project. We would also like to thank the Systems Biology Community at the University of Virginia for their invaluable feedback on the data analysis and the manuscript figures. The results shown here are in part based upon data generated by the TCGA Research Network: https://www.cancer.gov/tcga. Research reported in this publication was supported by the National Institute Of General Medical Sciences of the National Institutes of Health under Award Number R35GM138127 and by the National Science Foundation Graduate Research Fellowship under Grant No. 1842490. The content is solely the responsibility of the authors and does not necessarily represent the official views of the National Institutes of Health or the National Science Foundation.

## Declaration of generative AI and AI-assisted technologies

During preparation of this work, we used Claude.ai sonnet to identify grammatical errors and increase clarity of the manuscript, with a focus on making the objective of PTM-POSE and it’s utility more clear. After using this tool, we reviewed and edited the content as needed and take full responsibility for the content of the publication.

## Supporting Information

Supplementary Figure 1

Supplementary Figure 2

Supplementary Figure 3

Supplementary Figure 4

Supplementary Figure 5

Supplementary Figure 6

Supplementary Figure 7

Supplementary Figure 8

Supplementary Figure 9

Supplementary Figure 10

Supplementary Figure 11

Supplementary Figure 12

Supplementary Figure 13

Supplementary Figure 14

Supplementary Figure 15

Supplementary Figure 16

Supplementary Figure 17

Supplementary Figure 18

Supplementary Figure 19

Supplementary Figure 20

Supplementary Figure 21

Supplementary Table 1

Supplementary Table 2

Supplementary Table 3

Supplementary Table 4

