## Supplementary Information for "Systematic analysis of the effects of splicing on the diversity of post-translational modifications in protein isoforms using PTM-POSE"

### This file includes:

**Supplementary Figure 1**: General framework for enzyme-mediated, reversible post-translational modifications.
**Supplementary Figure 2**: Distribution of documented PTMs in canonical and alternative UniProt isoforms.
**Supplementary Figure 3**: Toy example of how PTMs are mapped and projected onto alternative transcripts.
**Supplementary Figure 4**: Impact of restricting analysis to PTMs with more evidence, either based on the number of studies observed or the number of different databases in which the PTM is found.
**Supplementary Figure 5**: Comparison of PTM density between constitutive and non-constitutive exons.
**Supplementary Figure 6**: Constitutive rate for all classes of post-translational modifications, if they have at least one documented PTM in the human phosphoproteome.
**Supplementary Figure 7**: Relationship between the presence of PTM in domain and its constitutive rate.
**Supplementary Figure 8**: Consitutive rate for subtypes within the classes in Supplementary Figure 6.
**Supplementary Figure 9**: Impact of using different criteria to define functionally relevant transcripts and PTMs on PTM constitutive rate and the number of isoforms per protein.
**Supplementary Figure 10**: Molecular functions and biological processes enriched within either constitutive or non-constitutive PTM sites.
**Supplementary Figure 11**: Impact of different splice events on PTMs.
**Supplementary Figure 12**: Overlap of significantly spliced phosphorylation sites with phosphorylation measured by mass spectrometry in KRAS-induced lung cells.
**Supplementary Figure 13**: Additional context to flanking sequences altered by splice events not discussed in the main text Figure 3.
**Supplementary Figure 14**: Summary of PTMs impacted by ESRP1-related splicing, including breakdown by modification type and comparison to ESRP1-knockdown experiments.
**Supplementary Figure 15**: Enriched gene processes, functions and processes for genes regulated by ESRP1 expression.
**Supplementary Figure 16**: Functional associations of PTMs impacted by ESRP1-correlated splicing.
**Supplementary Figure 17**: PTM-associated interactions most impacted by ESRP1-correlated splicing.
**Supplementary Figure 18**: Impacts of altered flanking sequences on interaction motifs, including impact on canonical 14-3-3 protein binding.
**Supplementary Figure 19**: Substrates of kinases that are enriched amongst ESRP1-regulated phosphorylation sites, in particular those of SGK1.
**Supplementary Figure 20**: Comparison of differential inclusion of phosphorylation sites and measured phosphorylation abundance in CPGEA prostate cancer patients.
**Supplementary Figure 21**: Relationship between kinase activity, substrate inclusion, and ESRP1 expression in the CPGEA prostate cancer cohort.

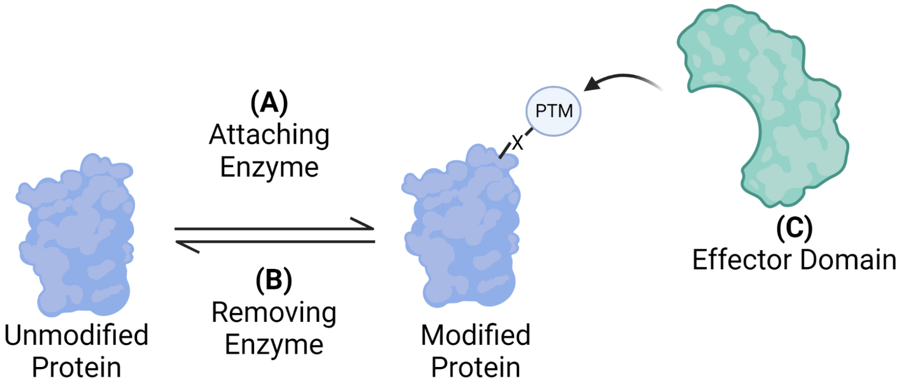


| Modification | Primary Residues (>100 Instances) | A) Adding Enzyme(s) | B) Removing Enzyme(s) | C) Effector Domains |
|---|---|---|---|---|
| Phosphorylation | S, T, Y | Kinase | Phosphatase | - Phosphoserines/threonines: 14-3-3, WW<br>- Phosphotyrosines: SH2, PTB |
| Ubiquitination | K | E1/E2/E3 Cascade | Deubiquitinases | - Ubiquitin-binding domains |
| Acetylation | K, M, S, A, T | Acetyltransferases | Deacetylase | - Acetyllysine: bromodomain |
| Methylation | R, K | Protein methyltransferases (PRMT) | Demethylase | - Methyllysine: Tudor, Chromo, Agenet, PWWP, WD40, MBT, Ank, BAH<br>- Methylarginine: Tudor |

**Supplementary Figure 1. Framework for most enzyme mediated post-translational modifications.** Most post-translational modifications are the result of reversible chemical reactions, with one class of enzymes responsible for adding the modification and a separate class of enzymes responsible for its removal. Further, many post-translational modifications are recognized by specific "reader" domains, and thereby help to facilitate protein-protein interactions. We provide examples of this framework for the four most common modification types: phosphorylation, ubiquitination, aceytlation, and methylation. Created with Biorender.com.

**A**

| Source | Data Type | Number of Instances |
| --- | --- | --- |
| Ensembl | Genes | 19,703 |
| | Transcripts | 89,110 |
| | Alternative Isoforms | 51,133 |
| UniProtKB | Proteins | 18,973 |
| | Alternative Isoforms | 32,602 |
| ProteomeScout/PhosphoSitePlus | Post Translational Modifications (PTMs) | 395,822 |
| | Genes/Proteins with PTMs | 17,663 (93.1%) |
| | UniProtKB Alternative Isoforms with annotated PTMs | 2124 (15.3%) |

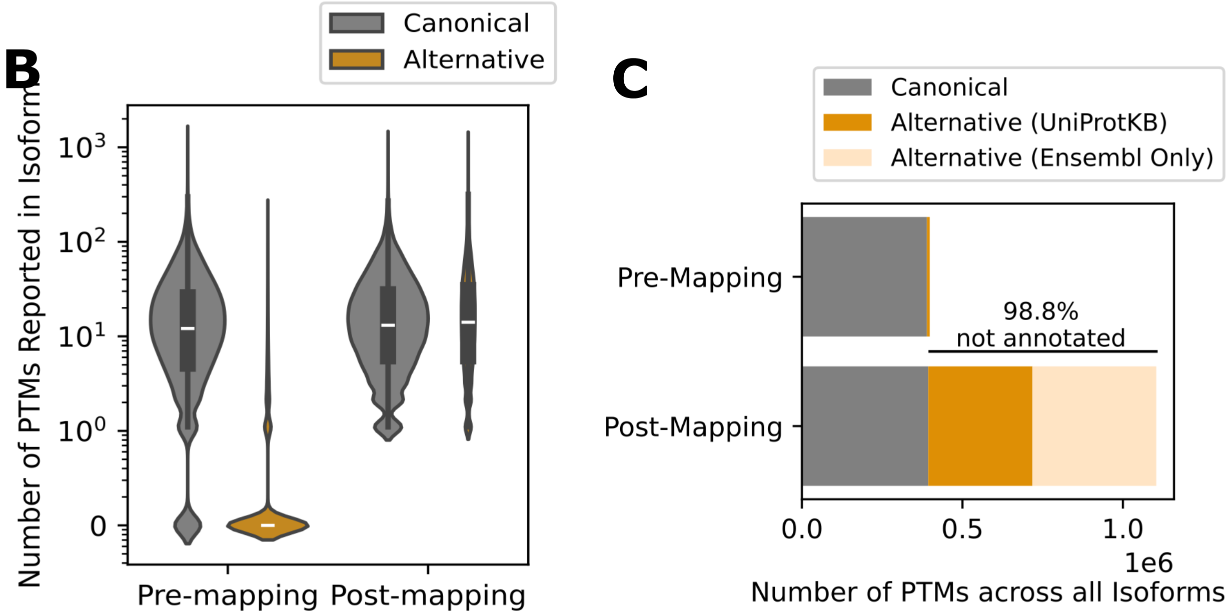


**Supplementary Figure 2. PTMs mapping and projection pipeline expands the available information of PTMs in alternative isoforms.** To expand the available information on PTMs in alternative isoforms, we pulled data from Ensembl [1], UniProtKB [2], ProteomeScout [3], and PhosphoSitePlus [4]. **A)** Total number of transcripts, proteins, isoforms, and post translational modifications utilized in the mapping/projection pipelin. We also highlight the fraction of proteins and alternative isoforms with annotated PTMs. In a 84.7% of cases, the alternative isoforms did not have any documented PTMs. **B)** Number of PTMs annotated for individual protein isoforms in ProteomeScout and PhosphoSitePlus (Pre-mapping), and the number for those same proteins after employing the mapping/projection pipeline. Proteins are separated by whether they are the canonical or alternative isoform in UniProt. The y-axis is on a log10 scale. **C)** Total number of PTMs identified in canonical and alternative isoforms before and after mapping/projection. Post-mapping data includes additional protein isoforms extracted from Ensembl, in addition to those already annotated in UniProt. We found that 98.8% of the PTMs we identified by mapping were not previously annotated ($\frac{1-Number\ of\ PTMs\ annotated\ in\ UniProtKB\ Isoforms}{Number\ of\ PTMs\ identified\ by\ Mapping}$).

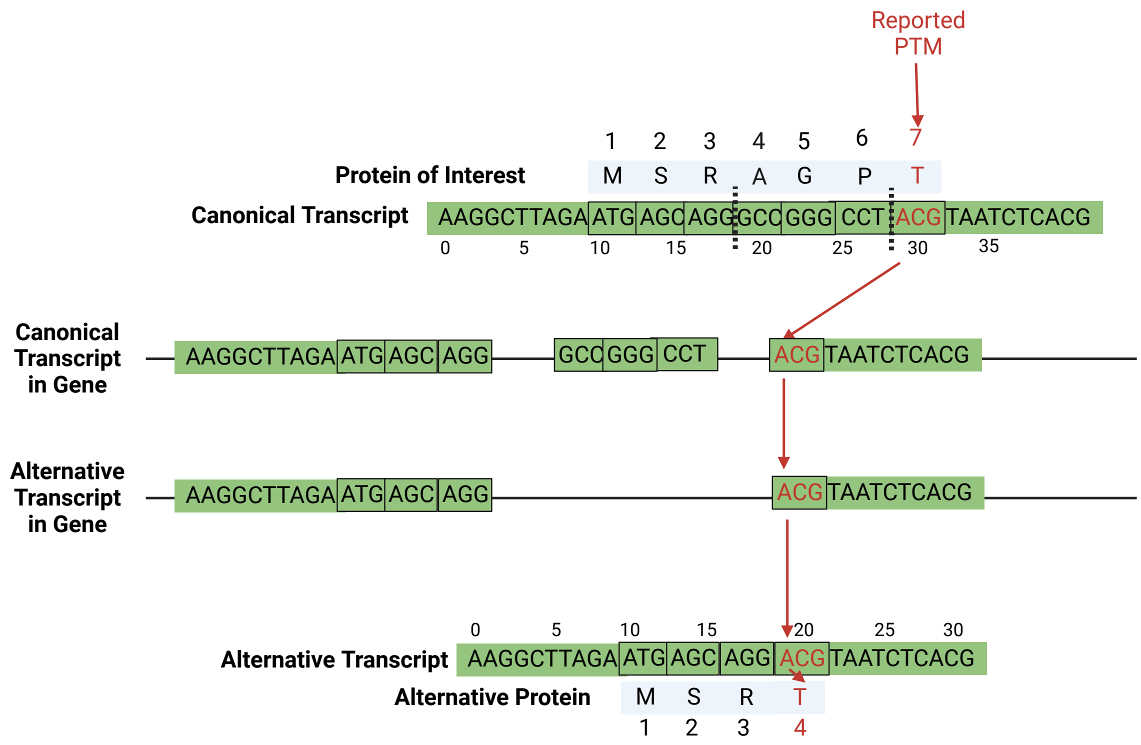


**Supplementary Figure 3. Toy example of the mapping/projection of PTMs onto alternative transcripts.** Here, we are illustrating how documented PTMs are mapped onto the genome and projected onto alternative transcripts, using a small mock protein/transcript as an example. **Phase 1, Mapping:** First, we identify a PTM and its location in the canonical isoform of the protein of interest, using information from ProteomeScout [3] and PhosphoSitePlus [4]. Using the location of the coding sequence in the transcript, we can then find the codon responsible for the PTM of interest, as well as the exon that the PTM can be found in (denoted by the dotted lines in the canonical transcript). Once the exon containing the PTM has been identified, we can use the locaiton of the exon in the genome to map the PTM to its genomic location. **Phase 2, Projection:** With the genomic location of the PTM, we can now look at the alternative transcripts produced by the gene and check if the genomic location of the PTM is utilized in the alternative transcript (based on genomic locations of exons in the alternative transcript). If it is not, we deem the PTM missing from the alternative isoform. If it is, we use the location of the coding sequence in the alternative transcript to project the PTM onto the alternative isoform. Finally, we perform a couple of final checks: 1) ensure projected site is still in a coding region, 2) make sure no frame shifts have disrupted amino acid sequence, 3) ensure that residue is unchanged and projection was accurate. Created with Biorender.com.

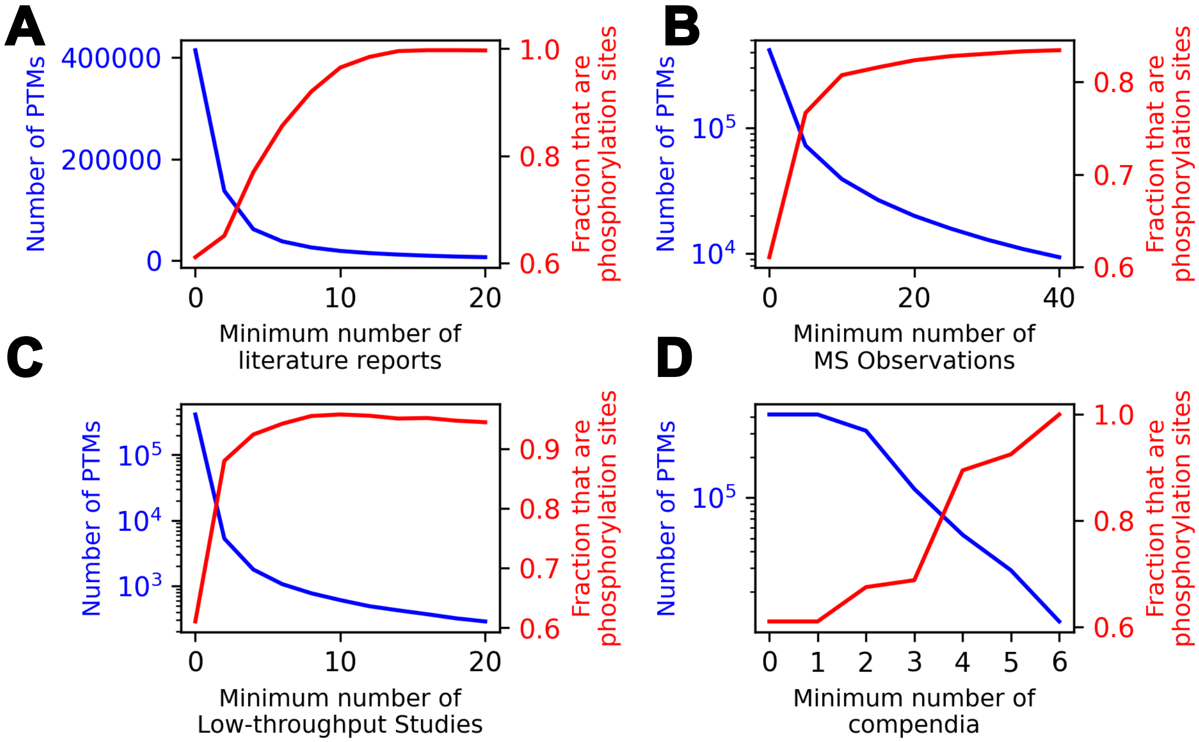


**Supplementary Figure 4. Impact of filtering PTM sites with less experimental evidence.** PTM-POSE provides options for filtering PTMs based on the degree of evidence supporting the PTM based on data from PhosphoSitePlus [4] and ProteomeScout [3]. Each plot shows the number of PTMs available for analysis (left axis, blue) or the fraction of available PTMs that are phosphorylation sites (right axis, red). PTMs can be filtered by: **A)** Number of literature reports (low throughput or by mass spectrometry)**B)** Number of mass spectrometry observations (in literature and by PhosphoSitePlus) **C)** Number of low-throughput literature studies **D)** Number of databases/compendia the PTM is recorded in. Possible databases in this analysis include ProteomeScout [3], PhosphoSitePlus [4], UniProt [2], dbPTM [5], HPRD [6], PhosphoELM [7]. As can be seen by each plot, while more stringent filters will focus on well-studied sites, it also restricts the overall number of PTMs available for analysis and results in a larger skew towards phosphorylation sites.

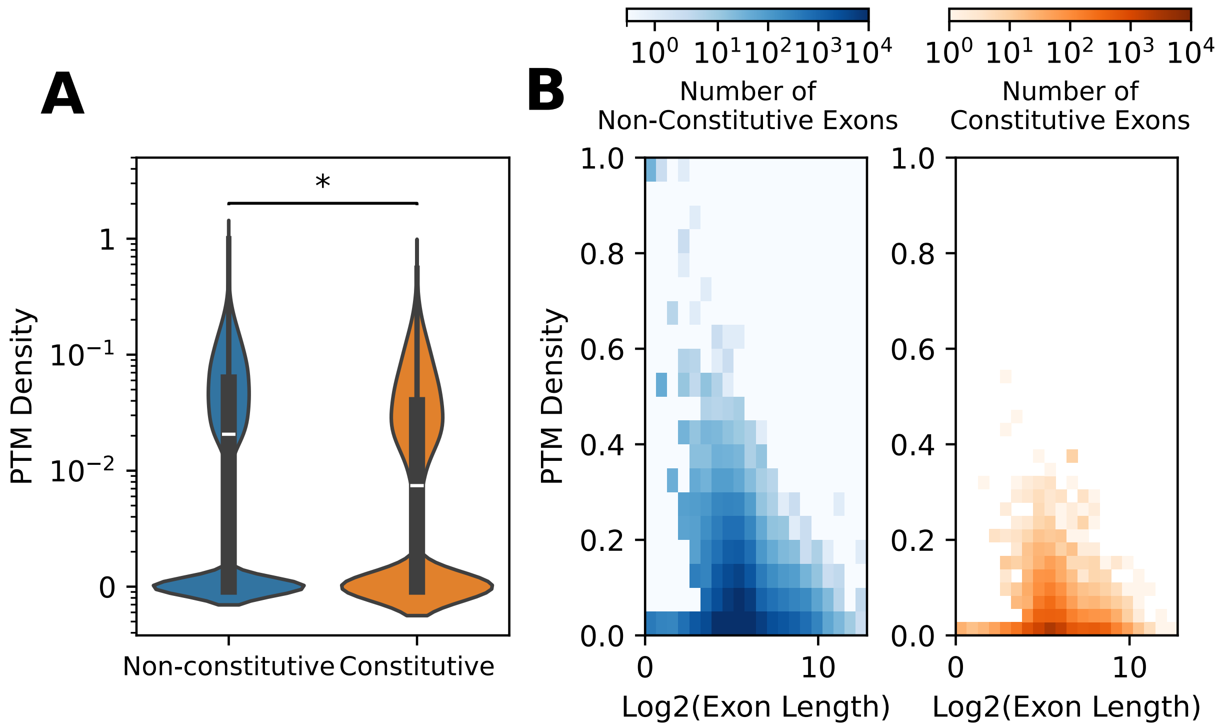


**Supplementary Figure 5. Constitutive exons exhibit lower PTM density than non-constitutive exons.** Comparison of density of PTM sites in constitutive exons vs. non-constitutive exons, as defined by Ensembl. We calculated PTM density as the number of PTM sites mapped to an exon divided by the number of total residues coded for by the exon. **A)** Violin plot comparing density of PTMs between constitutive and non-constitutive exons. Significance was assessed using a two-tailed Mann Whitney U test (*: $p < 0.05$). The y-axis is on a log10 scale. **B)** Histogram showing the relationship between PTM density and exon length (by number of residues) for both non-constitutive (blue) and constitutive (orange) exons. Colorbars are on a log-scale and illustrate the number of exons without that PTM density and exon length. Maximum value shown is 1000 exons.

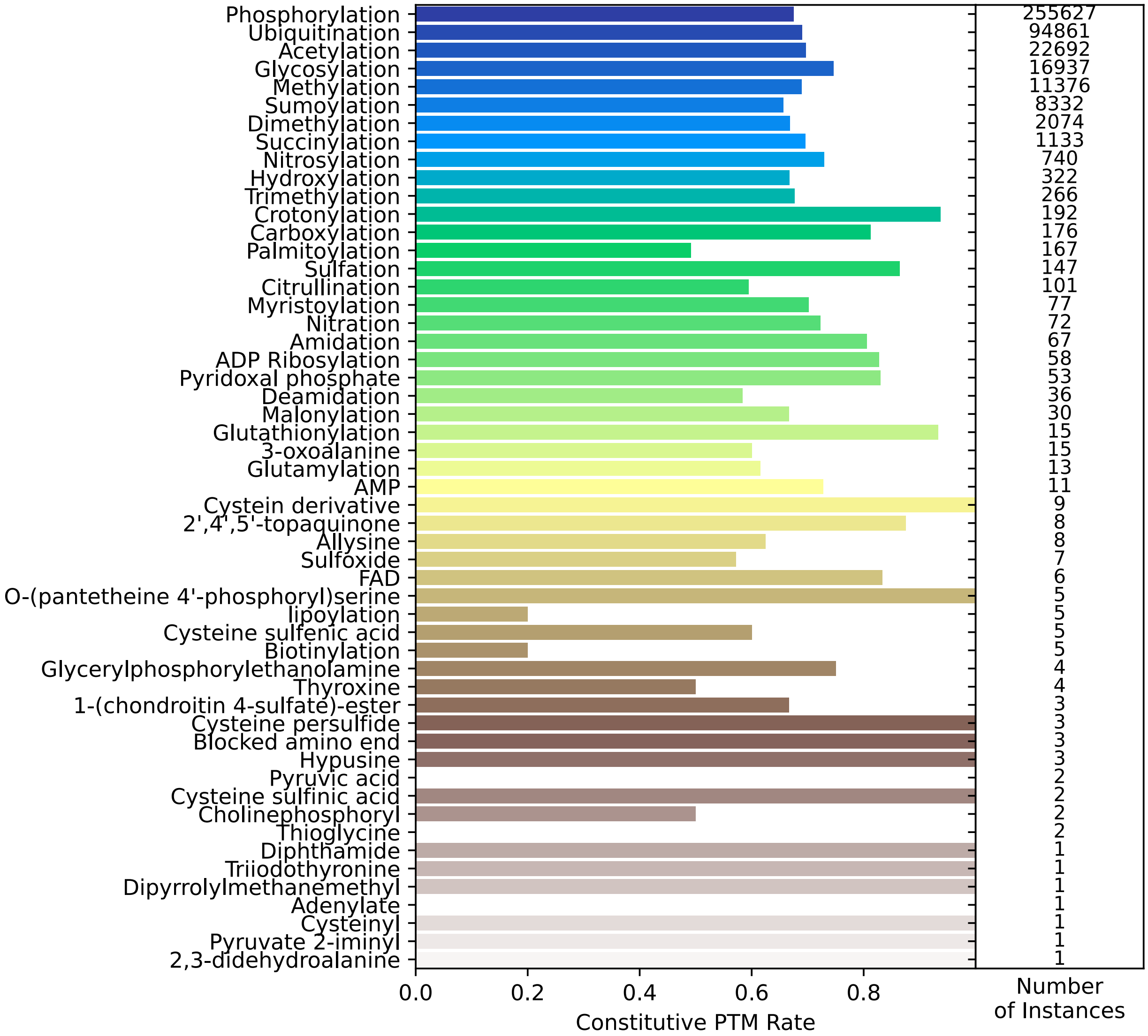


**Supplementary Figure 6. Constitutive PTM rate across different modification types.** The fraction of PTMs that are found across all alternative isoforms of a given protein (constitutive rate), broken down by broad modification class. This is an extended version of Figure 2B, but without the comparison against a null model due to computational time constraints.

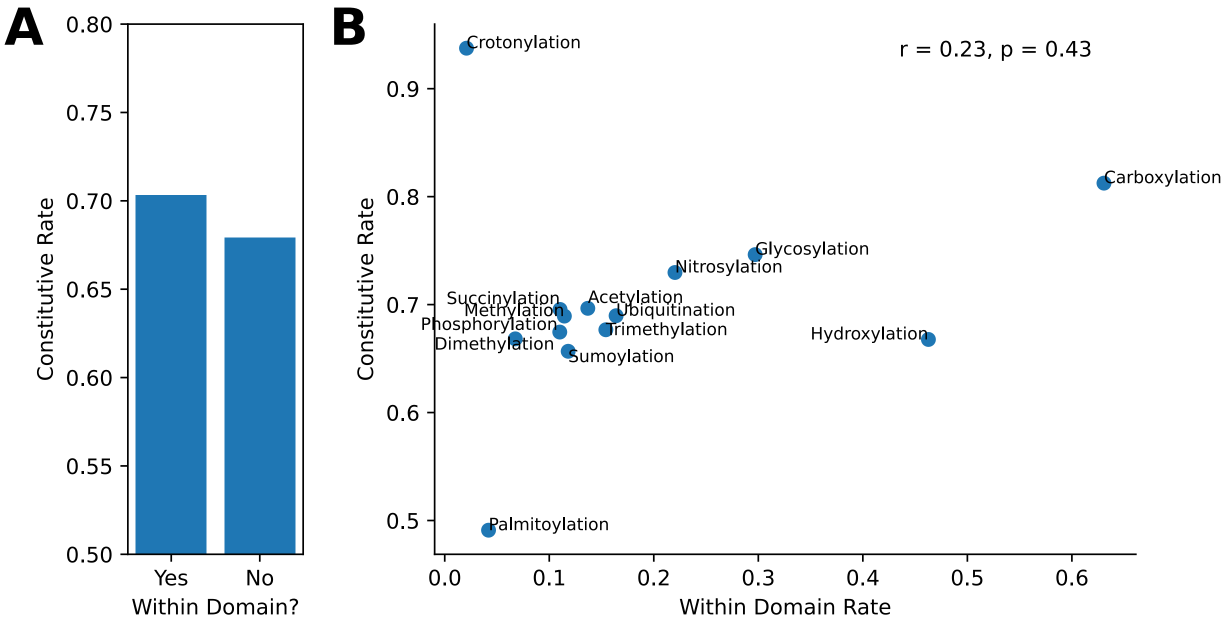


**Supplementary Figure 7. PTMs found within conserved structural domains are slightly less likely to be regulated by splicing.** Given that it has been previously illustrated that disordered regions are more likely to exist in spliced exons than structural regions, we were curious to know whether the constitutive rate of PTMs found within and outside of conserved structural domains exhibited differences in constitutive rate (fraction of PTMs that are found across all isoforms of the gene).**A)** Overall constitutive rate for PTMs found within domains versus those found outside of domains. **B)** Relationship between the fraction of a modification type found within a domain and its constitutive rate. Although increased presence in domains increases constitutive rate for some PTMs (i.e. glycosylation), we did not find a consistent relationship that explains constitutive rate across modification types.

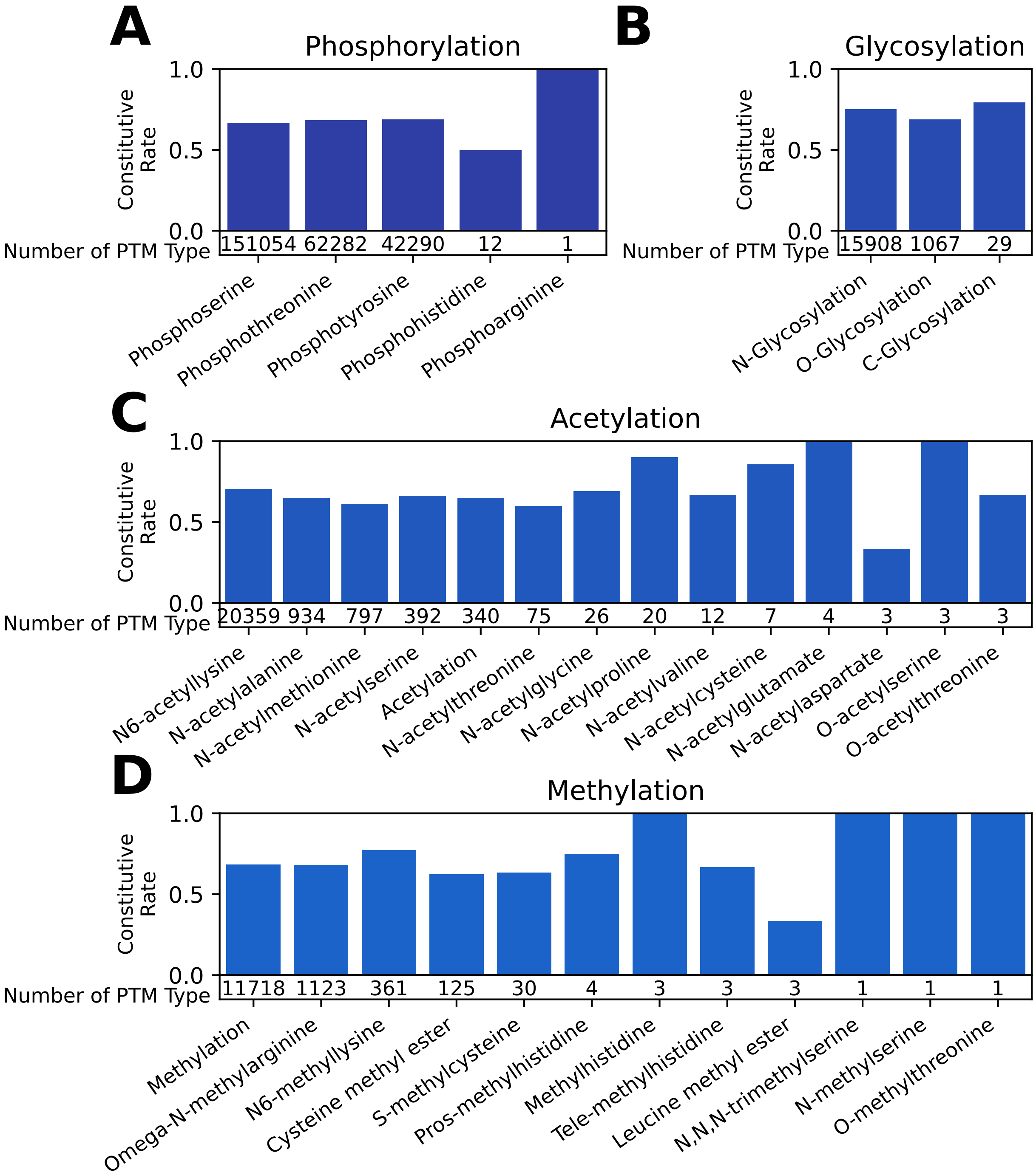


**Supplementary Figure 8. Constitutive PTM rate across different modification types** Constitititutive rates for modification subtypes. This is the same analysis as Supplementary Figure 6, but modifications have been further broken down into their subtypes. For example, for phosphorylation, we also looked at the differences between phosphoserines, phosphothreonines, and phosphotyrosines. (continued on the next page)

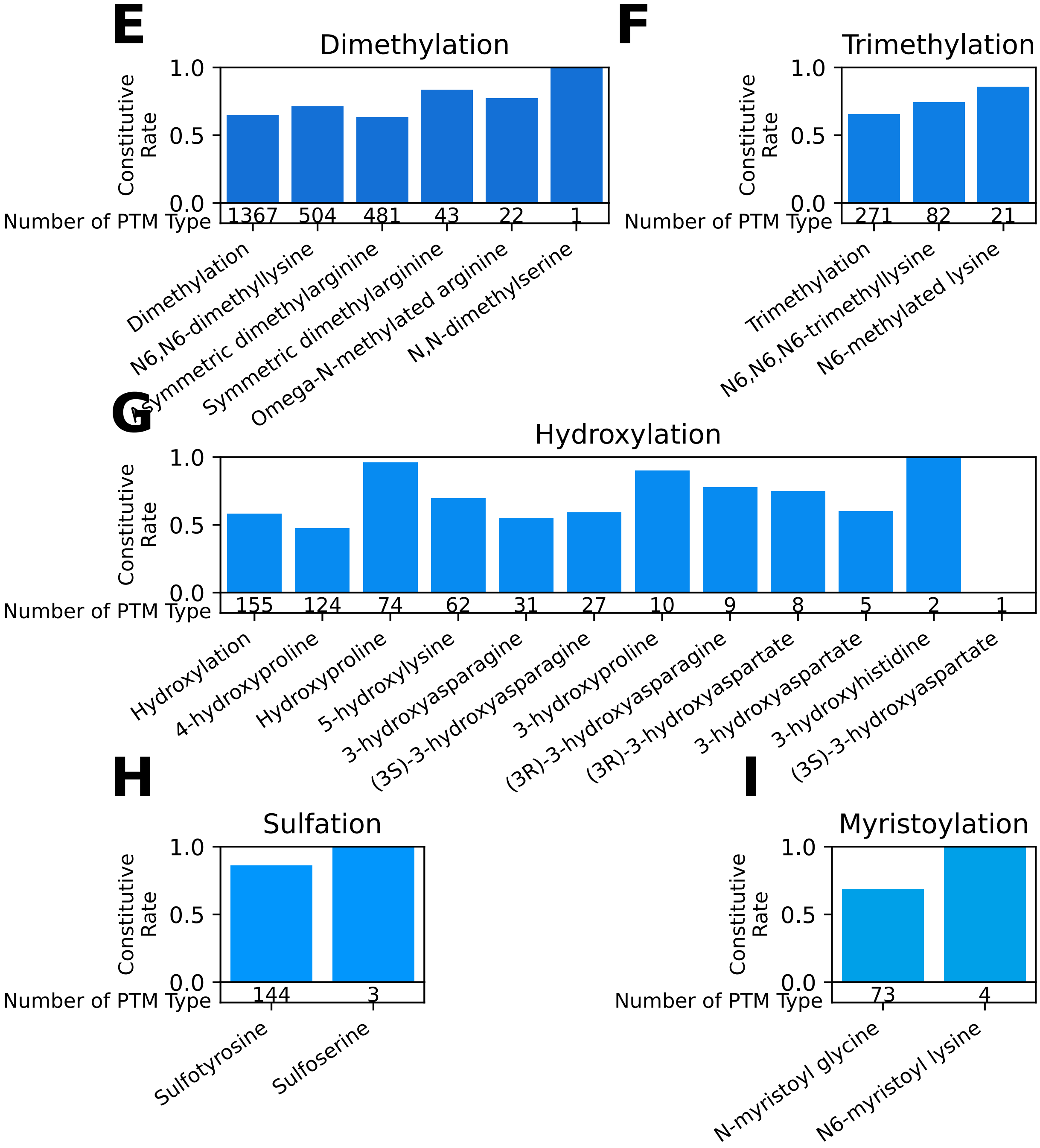


**Supplementary Figure 8. Constitutive rate for modification subtypes** Constititutive rates for modification subtypes. This is the same analysis as Supplementary Figure 6, but modifications have been further broken down into their subtypes. For example, for phosphorylation, we also looked at the differences between phosphoserines, phosphothreonines, and phosphotyrosines (Continued on next page)

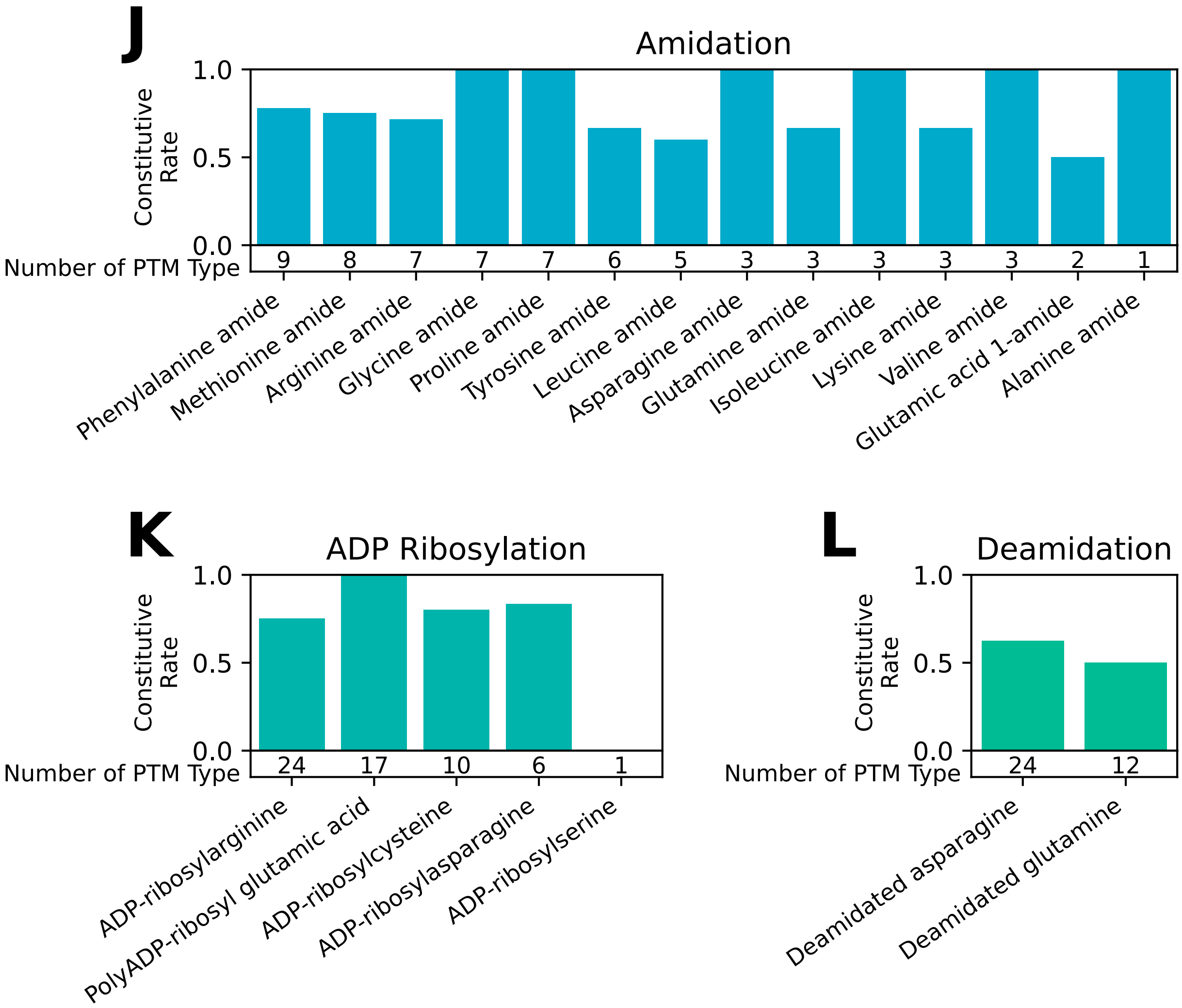


**Supplementary Figure 8. Constitutive rate for modification subtypes.** Constititutive rates for modification subtypes. This is the same analysis as Supplementary Figure 6, but modifications have been further broken down into their subtypes. For example, for phosphorylation, we also looked at the differences between phosphoserines, phosphothreonines, and phosphotyrosines

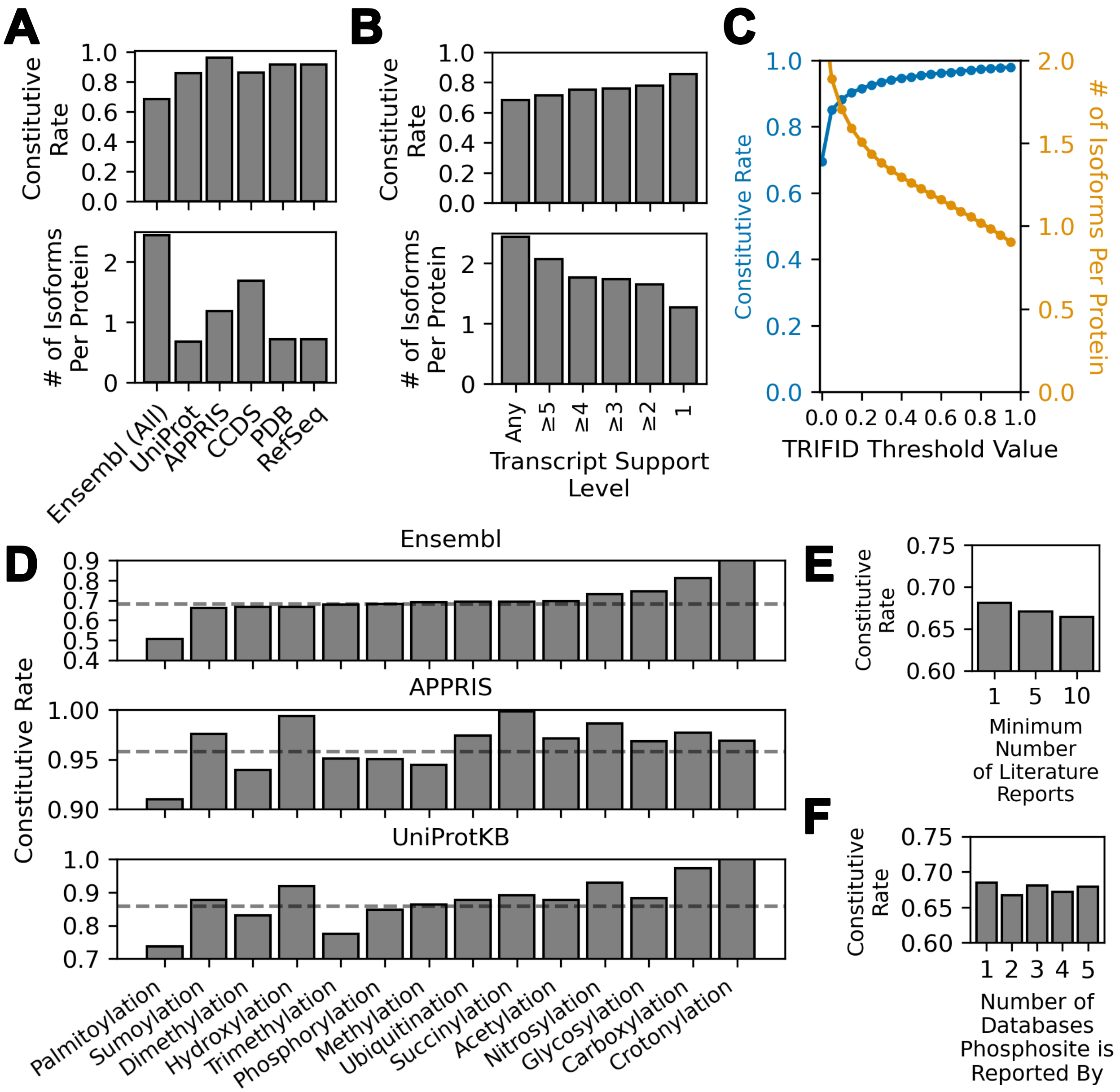


**Supplementary Figure 9. Constitutive PTM rate using different definitions of a functional transcript and PTMs.** We looked at the number of isoforms per protein and the constitutive rate (fraction of PTMs that are found in all isoforms of a given protein) after transcripts were filtered using various criteria. **A)** Filtering based on whether the transcript is found in various genomic or proteomic database, including: Consensus Coding Sequence Project (CCDS) [8], NCBI Reference Sequence (RefSeq) [9], APPRIS [10], UniProt [2], and Protein Data Bank (PDB) [11]. **B)** Filtering based on experimental transcript support level (TSL), as defined by Ensembl using mRNA and EST alignments [1]. TSL1 transcripts are the most strongly supported, while TSL5 transcripts have limited evidence. **C)** Filtering based on predicted functionality of each transcript using the TRIFID algorithm [12]. A TRIFID score of 1 indicates the most functional version of a gene, while 0 indicates lack of functionality. **D)** Modification-specific constitutive rates using Ensembl, UniProtKB, or APPRIS-defined transcripts. Modifications are sorted by the constitutive rates found when using all transcripts found in Ensembl. The data shown for Ensembl rates is identical to those found in Figure 2B of the main text. **E)** Overall constitutive rate if restricting assessed PTMs to those measured by mass spectrometry in at least 1, 5, or 10 different publications, based on data from PhosphoSitePlus [4]. **F)** Overall constitutive rates of PTMs found in 1, 2, 3, 4, or 5 different databases, based on information from ProteomeScout [3]. Databases assessed include PhosphoSitePlus [4], UniProt [2], dbPTM [5], PhosphoELM [7], and HPRD [6]

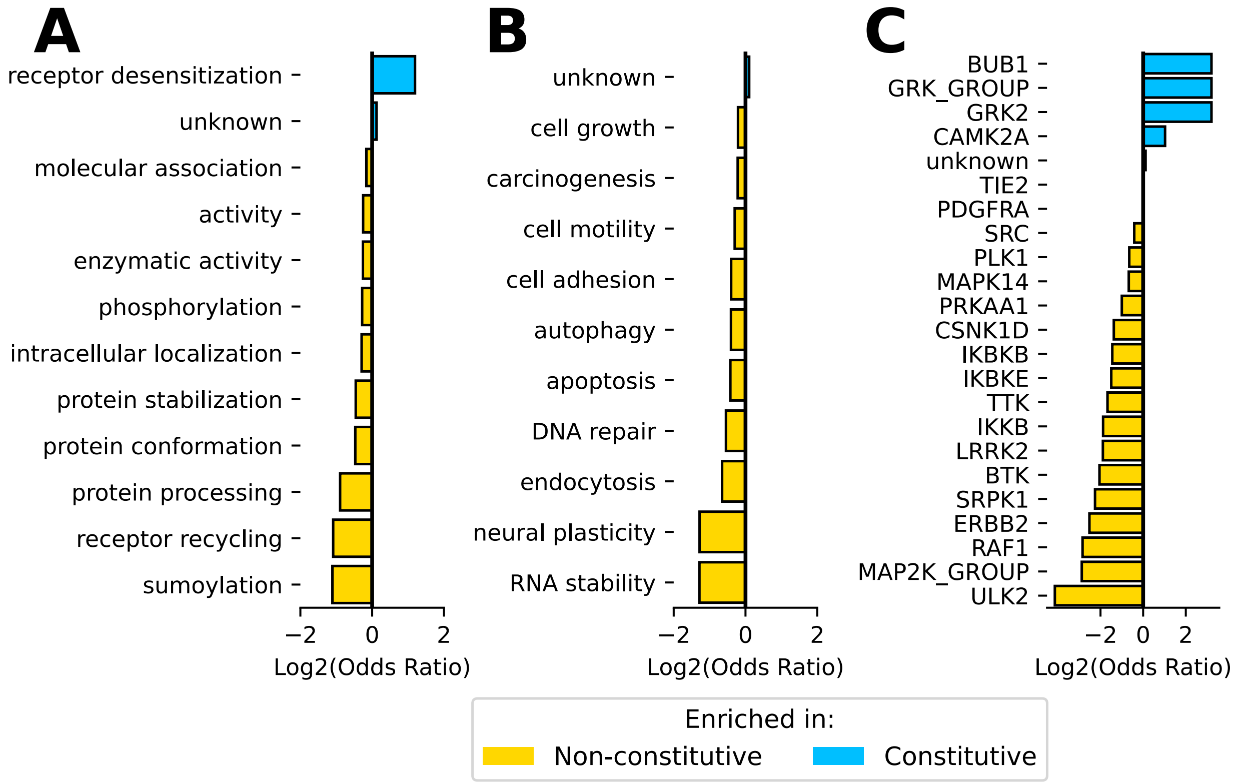


**Supplementary Figure 10. Splicing-controlled PTMs are important for many different molecular functions and biological processes.** To assess if PTMs associated with certain molecular functions or biological processes of PTMs are more or less likely to be maintained in an alternative isoform, we downloaded molecular function, biological process, or kinase-substrate annotations for individual PTMs from PhosphoSitePlus [4] and RegPhos [13]. For each annotation, a Fisher's Exact test was employed to identify whether it was enriched in either non-consititutive or constitutive PTM sites. P-values were corrected using Benjamini-Hochberg false positive correction [14]. We then compared odds ratios for **A)** enriched molecular functions, **B)** enriched biological processes, or **C)** enriched kinase-substrate associations ($p \leq 0.05$). PTMs with no known annotated molecular function or process are labeled as "Unknown".

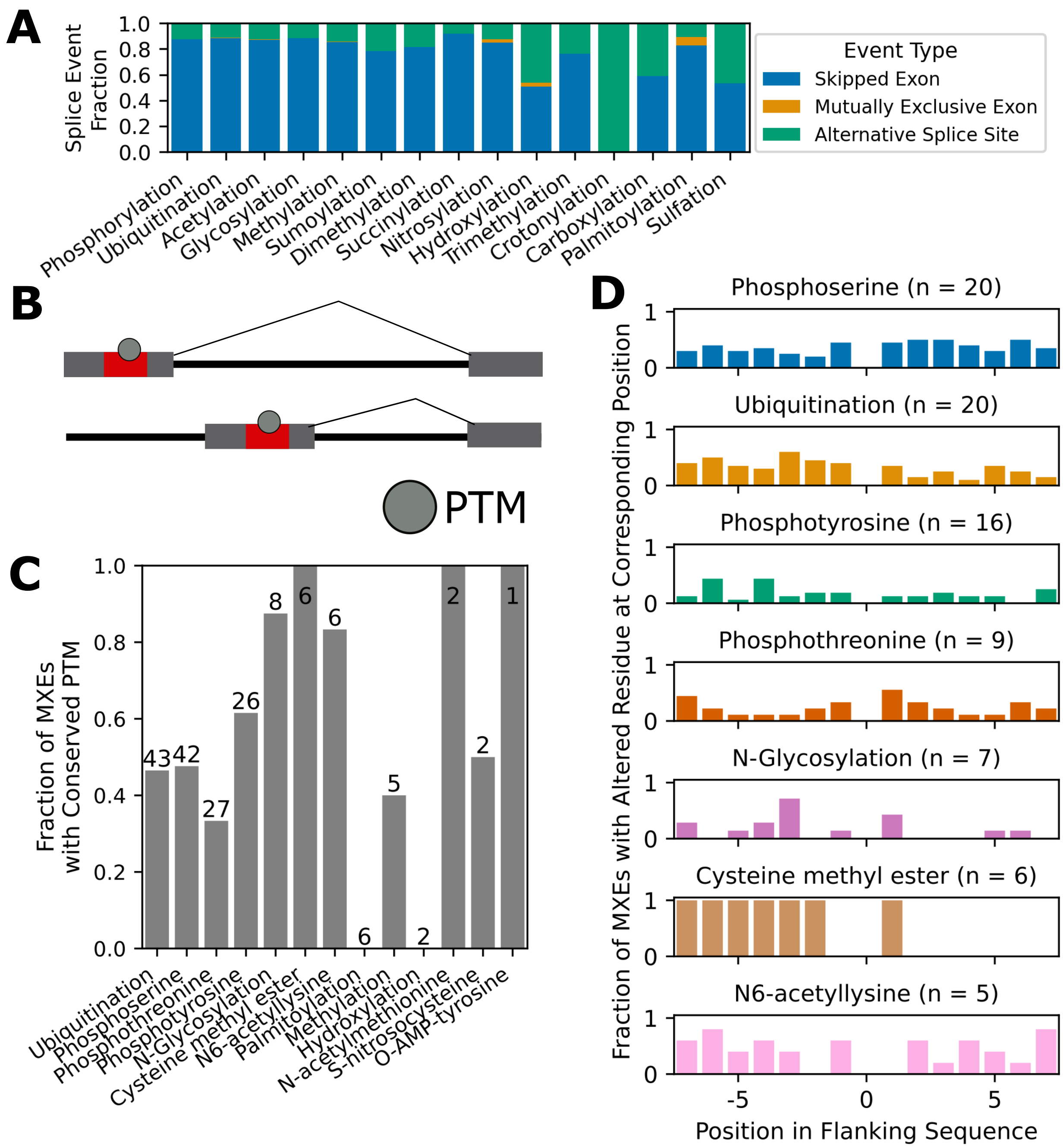


**Supplementary Figure 11. Role of different splice event types on regulating post-translational modifications. A)** Breakdown of the alternative splicing events that result in the loss of a PTM site relative to the canonical isoform, considering skipped exon events, alternative splice sites (ASS), and mutually exclusive exons (MXE). ASS are defined as described in Wang et al. and elsewhere [15]. MXEs were identified based on criteria from Pillmann et al. where MXEs are nearby the original exon, have exon lengths within 20 amino acids of each other, and have at least 15% sequence similarity [16]. **B)** Visualization of conservation of PTMs from mutually exclusive exons. **C)** Fraction of MXE events which conserve the PTM site found in the canonical isoform. Numbers above each bar indicate the number of that modification that was identified across all mutually exclusive exon events. **D)** For cases in which a PTM was found maintained in both MXEs, we asked which of the residues surrounding the PTM were altered. Considering the PTM site as the central point (indicated as 0), we calculated the number of times the surrounding residues were changed in the alternative exon, and reported the results as the fraction of instances in which that the residue was changed.

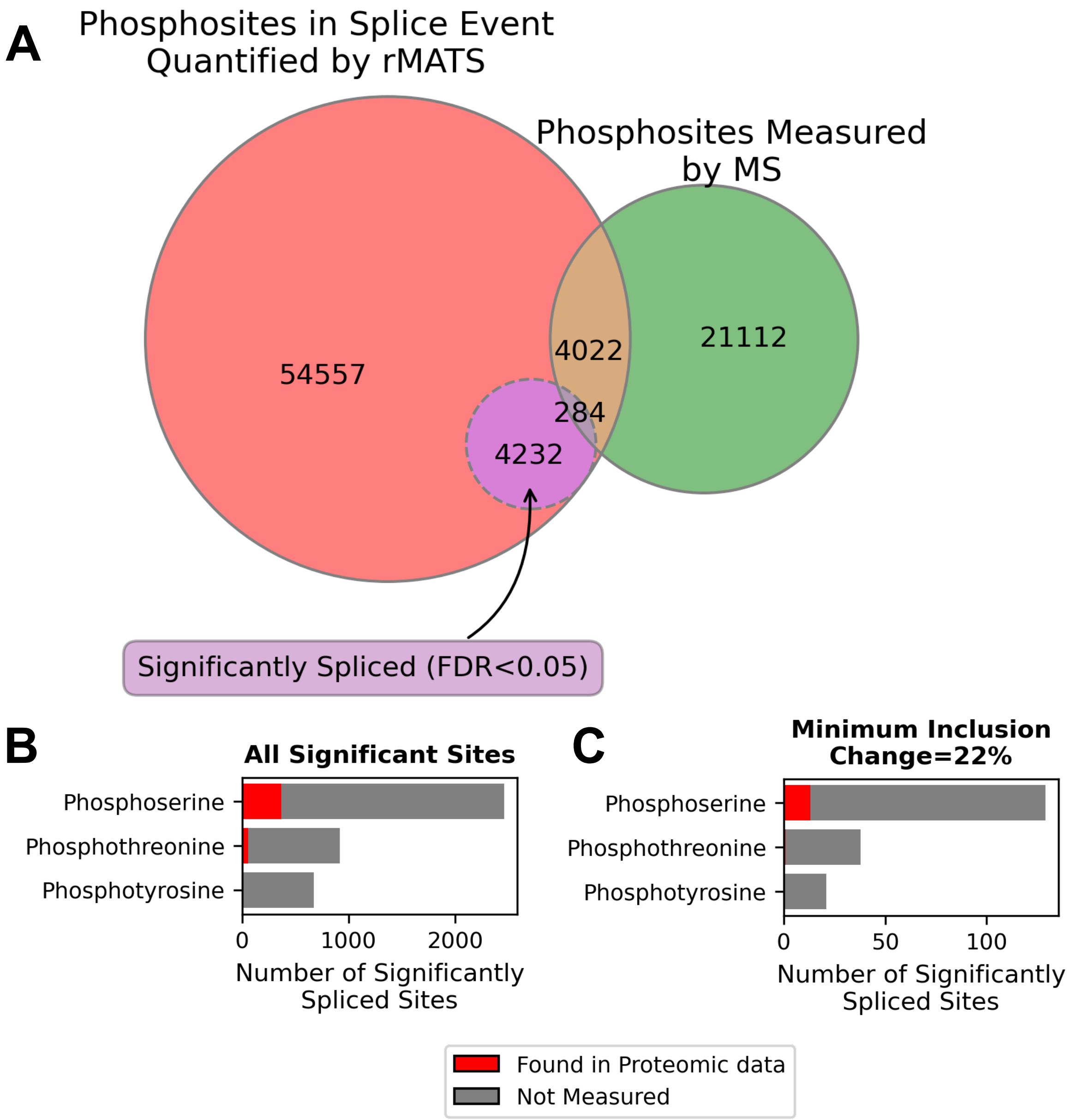


**Supplementary Figure 12. Overlap of significantly spliced phosphorylation sites with phosphorylation measured by mass spectrometry in KRAS-induced lung cells.** To compare differential inclusion of phosphorylation sites and measured phosphorylation, we identified phosphorylation sites that were significantly spliced after wild-type KRAS expression in AALE cell lines by applying rMATS [17] and PTM-POSE to RNA-sequencing data obtained by Lo et al. [18]. We then identified which of the spliced sites were also found in mass-spectrometry based phosphoproteomic data of the same cell lines. **A)** Overlap between sites identified from splicing analysis and phosphoproteomic measurement by MS, looking at all sites quantified by rMATS/PTM-POSE, as well as the ones with significant changes in inclusion. **B)** Number of each phosphorylation type (serine, threonine, or tyrosine) found to be significantly spliced in WT KRAS-transduced cells, including which sites were also measured by mass spectrometry (in red). **C)** Same as Panel B, but only including sites with at least 22% change in inclusion.

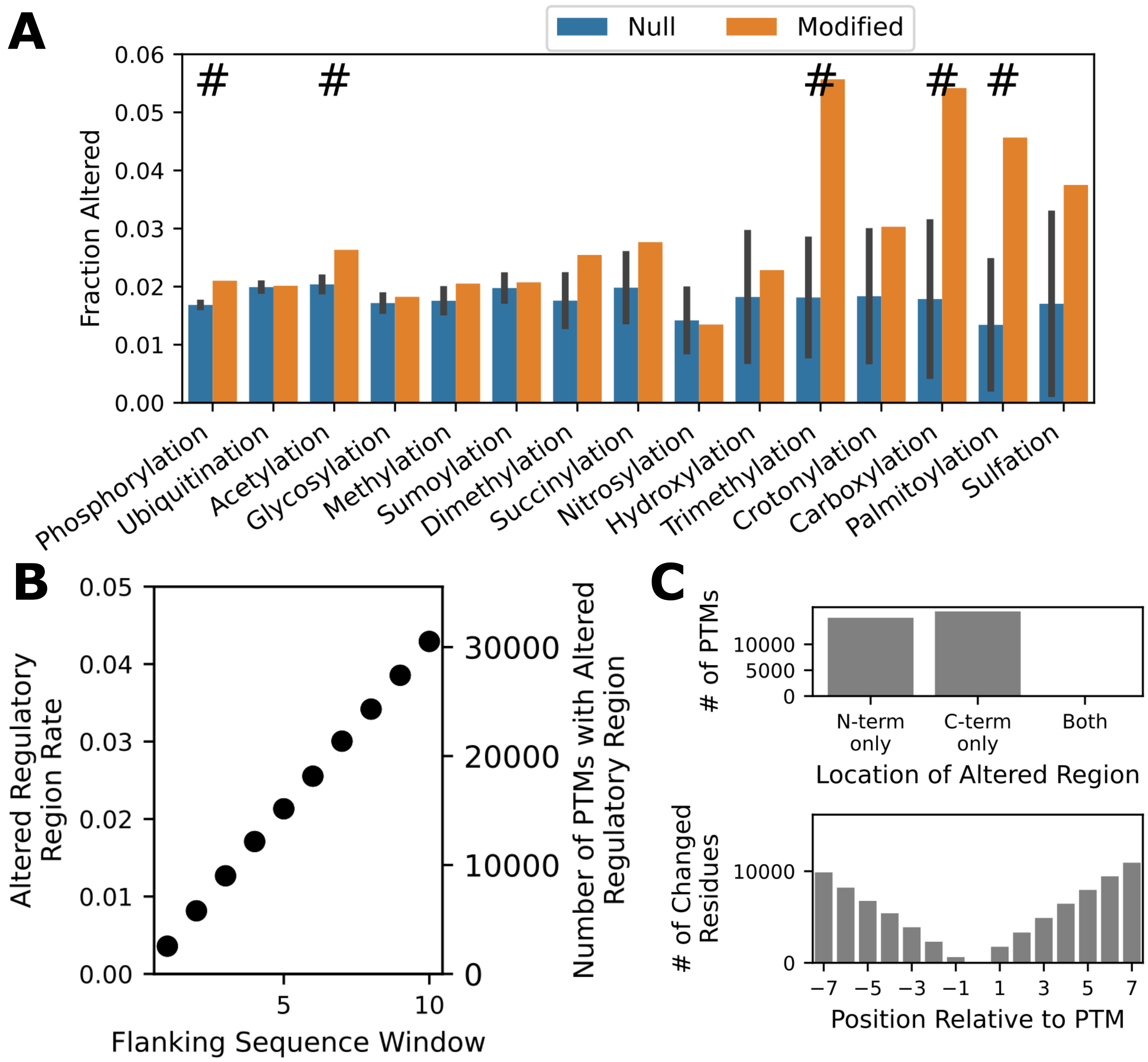


**Supplementary Figure 13. Exploring the prevalence of altered flanking sequences surrounding PTM sites.** Here, we have explored how often the residues surrounding the PTM site (termed flanking sequence) are altered and the location of these changes. **A)** Fraction of prospective PTM sites in alternative isoforms that have an altered regulatory sequence, considering the five residues on either side of the PTM site. We compared to a null model (as done in Figure 2B of the main text) which represents the rate expected if we were to randomly select residues from across the transcriptome. **B)** The overall fraction of prospective PTM sites with altered regulatory sequences when considering different number of residues surrounding the PTM site. As the regulatory window size increases, the fraction of altered PTMs also increases. **C)** Location of the residues that are altered when considering a regulatory sequence window of seven amino acids, either considering just the N-terminal side or C-terminal side (top figure), or the specific residue positions surrounding the PTM site at position 0 (bottom figure).

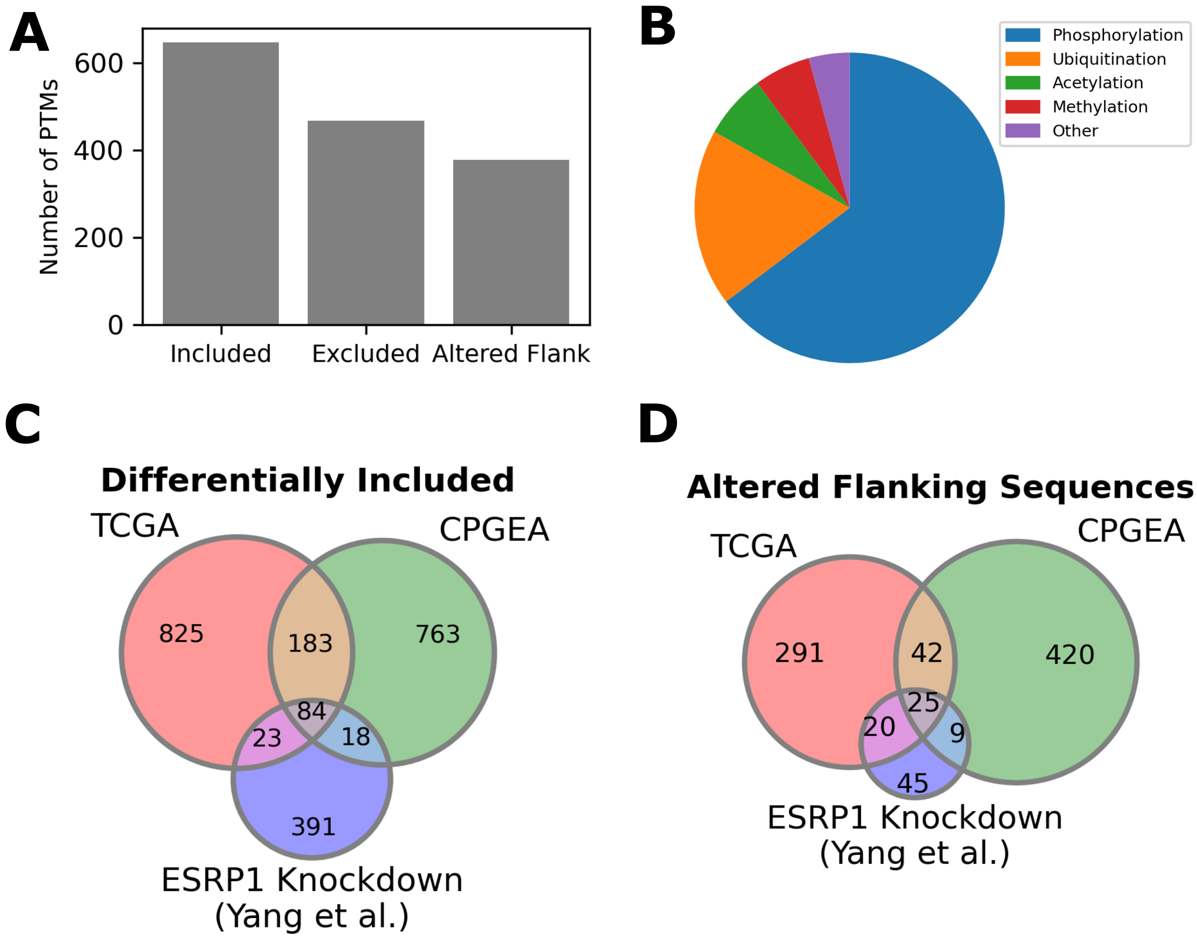


**Supplementary Figure 14. Summary of ESRP1-regulated PTM sites identified across different ESRP1-knockdown studies and prostate cancer cohorts.** In order to assess the role of ESRP1 in modulating PTM sites, we applied PTM-POSE to splice events quantified in either ESRP1-knockdown studies [19] or prostate cancer cohorts from The Cancer Genome Atlas (TCGA) or (CPGEA). Splice events were either quantified by rMATS (Yang et al. [19], CPGEA [20]) or SpliceSeq (TCGA [21]). Splice events and their corresponding PTMs related to ESRP1 expression were then extracted for each dataset. Further, PTMs in flanking exons nearby the splice junction were tested for altered flanking sequences based on whether the adjacent exon was included or excluded. **A)** Number of differentially spliced PTM sites in the TCGA dataset that are found predominantly in either ESRP1-low ($z \leq -1$) or ESRP1-high group ($z \geq 1$), or with altered flanking sequences. **B)** Distribution of modification types across differentially spliced PTMs in the TCGA cohort. **C)** Overlap between differentially included PTMs related to ESRP1 that could be identified within the TCGA prostate cancer cohort, from the CPGEA prostate cancer cohort, and from an ESRP1-knockdown experiment performed by Yang et al. [19]. **D)** Overlap between PTMs with altered flanking sequences due to ESRP1 expression in the same datasets as Panel C.

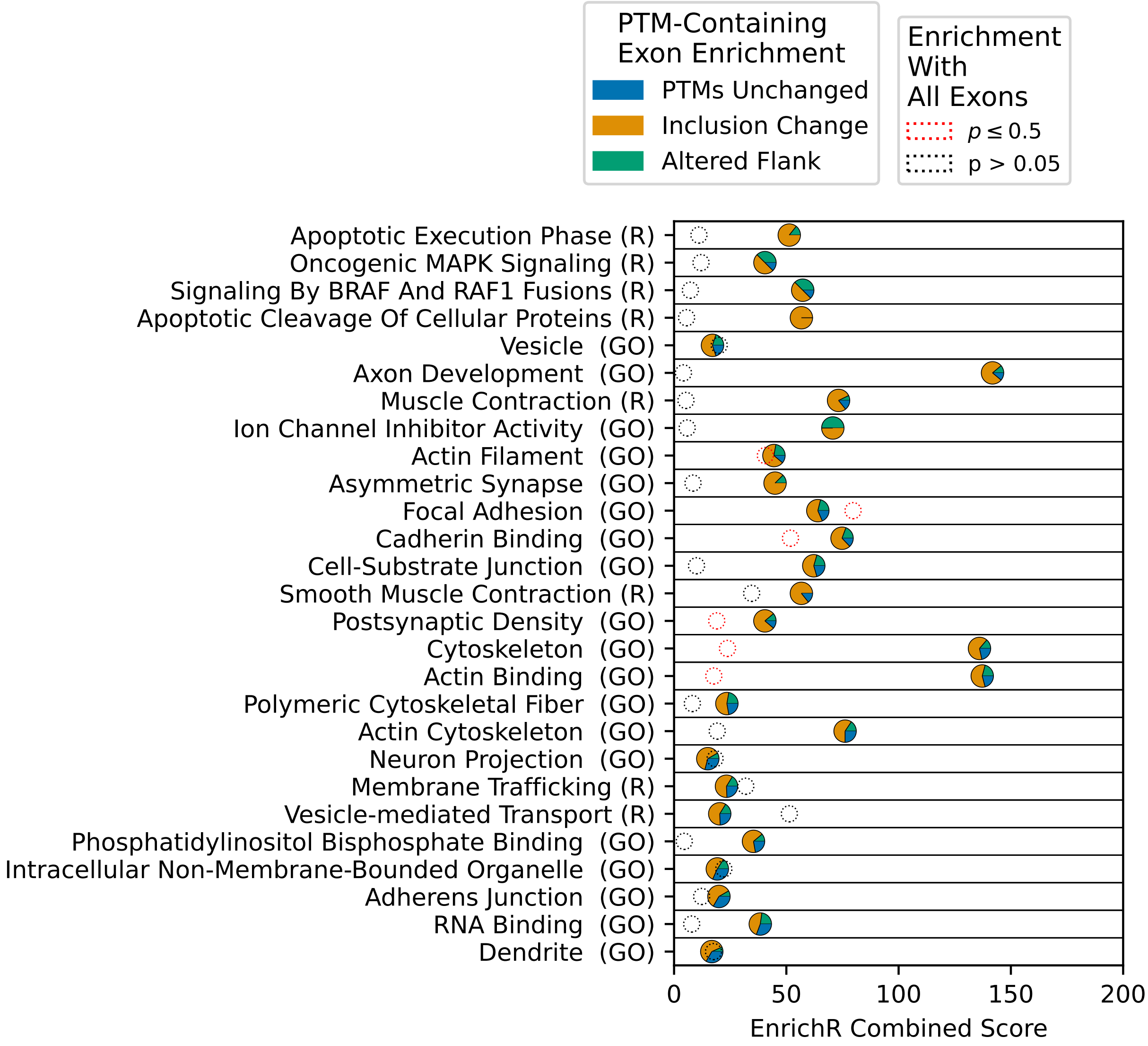


**Supplementary Figure 15. Genes with ESRP1-regulated PTMs are important to the actin cytoskeleton, apoptosis, and various other biological processes.** In order to better understand ESRP1 regulation in prostate cancer, we looked for enriched Gene Ontology terms (GO) [22] and Reactome pathways [23] among genes with differentially included PTMs or PTMs with altered flanking sequences. Enrichment was calculated with EnrichR [24], using the gseapy python package [25]. The background gene set consisted of all genes identified within the TCGA prostate cancer cohort. Two different enrichment scores are shown on the figure, one using all genes with differentially spliced exons (circles with dotted lines), one using only genes with spliced exons containing PTMs (pie charts). Each pathway/term shown on the figure was found to be significantly enriched among genes with spliced PTMs. Each pie chart (PTM-containing exon score) is colored to highlight the fraction of spliced genes associated with the term/pathway that belong to one of the three following groups: no PTMs impacted between isoforms, contains PTMs differentially included between isoforms, or only contains PTMs with altered flanking sequences. Terms are sorted based on the fraction of spliced genes that are associated with PTMs, with the most at the top and the least at the bottom.

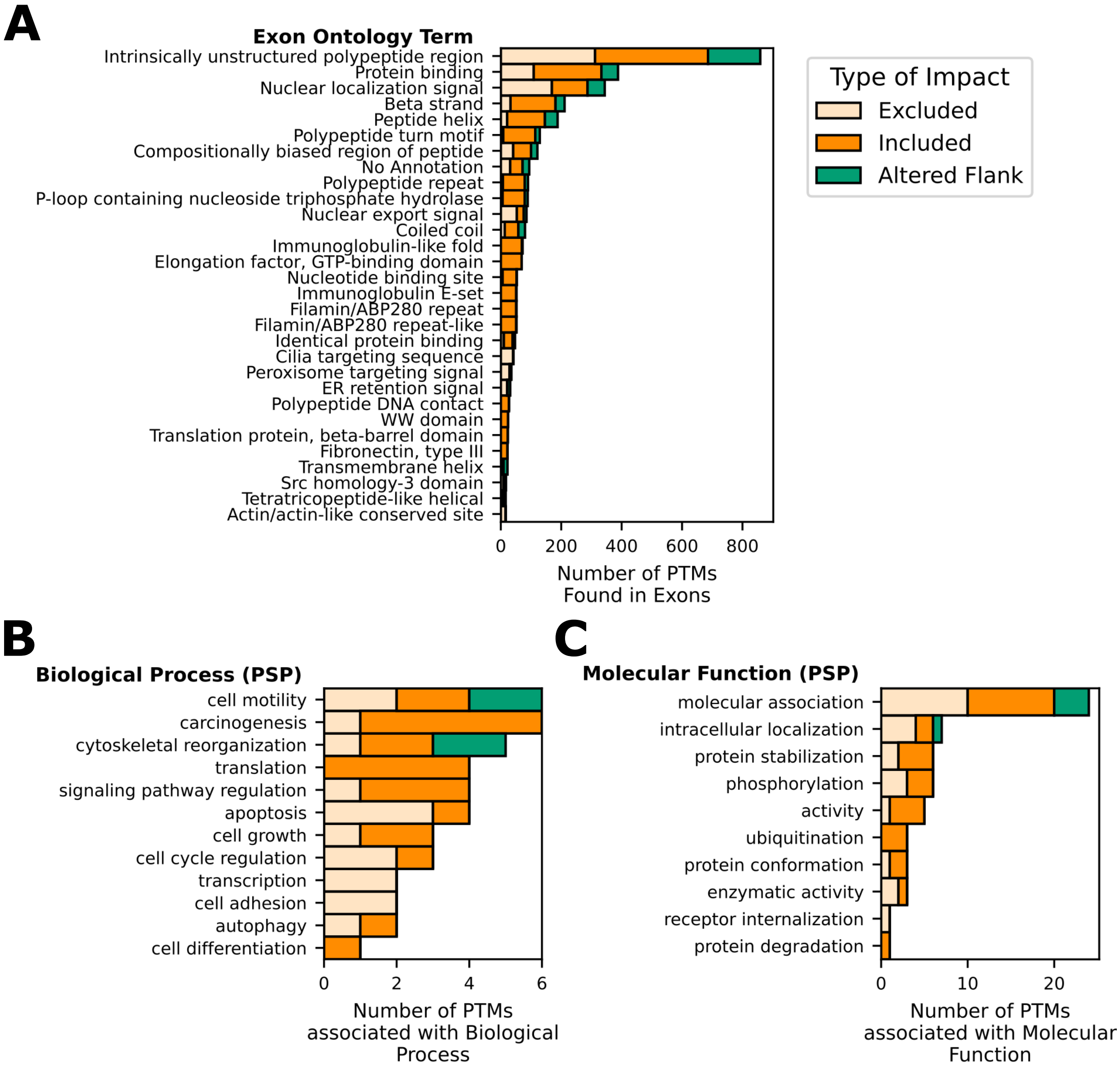


**Supplementary Figure 16. ESRP1-regulated PTM sites consist of modifications relevent to various functions and processes.** Number of PTMs and their functional associations which are either excluded, included, or contain altered flanking sequences in isoforms expressed with high levels of ESRP1 expression in prostate cancer ($z \geq 1$). PTM function was assessed based on **A)** Exon Ontology terms associated with the exon region containing the PTM [26], **B)** association with specific biological process based on annotations in PhosphoSitePlus [4]), or **C)** annotated molecular function in PhosphoSitePlus [4].

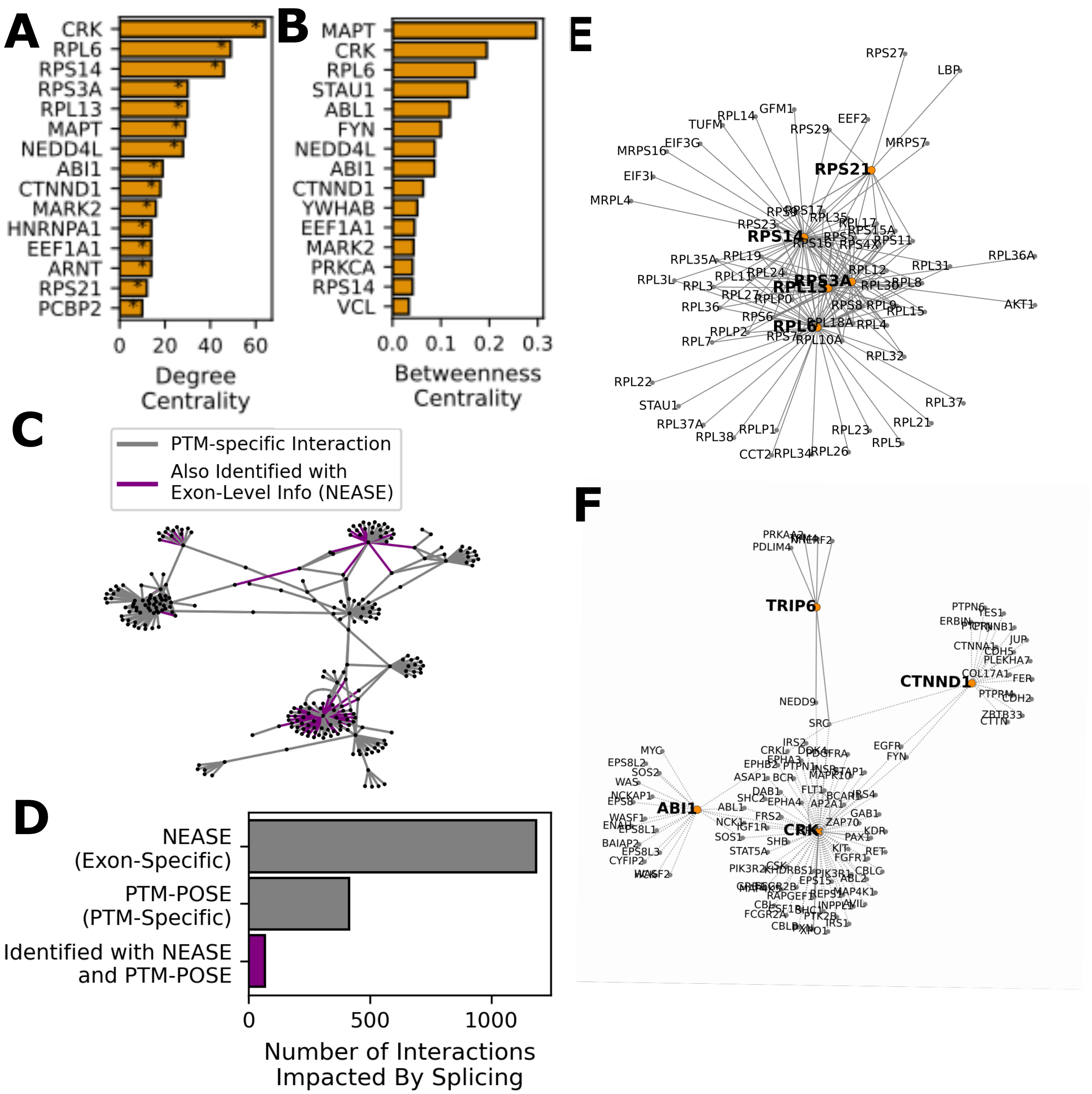


**Supplementary Figure 17. Altered PTM-associated interactions are centered around ribosomal proteins and those involved in cell signaling.** Based on PTM-associated interactions from PhosphoSitePlus [4], RegPhos [13], and PTMcode [27], we identified proteins with the most disrupted interactions due to differentially included PTMs due to ESRP1 expression in the TCGA prostate cancer dataset. **A)** Proteins within the highest degree in the altered interaction network. Overrepresentation of each protein was tested using a hypergeometric test, where the background network consisted of all annotated PTM-associated interactions. *:$p < 0.05$ **B)** Proteins with the highest betweenness centrality (indicator of proximity to all nodes in the network) in the altered interaction network. **C)** Network diagram of key PTM-driven interactions impacted by splicing, with edges colored in purple if the interaction was also identified by NEASE, which uses exon-level information such domain-domain interactions [28] **D)** Summary of the protein interactions impacted by splicing identified by NEASE [28], PTM-POSE, or both. **E)** Interactions for proteins associated with SRC/ABL signaling (CRK, ABI1, CTNND1, TRIP6). Proteins with spliced PTMs are indicated as orange nodes. **F)** Ribosomal protein associated increased with ESRP1 expression. Proteins with spliced PTMs are indicated as orange nodes.

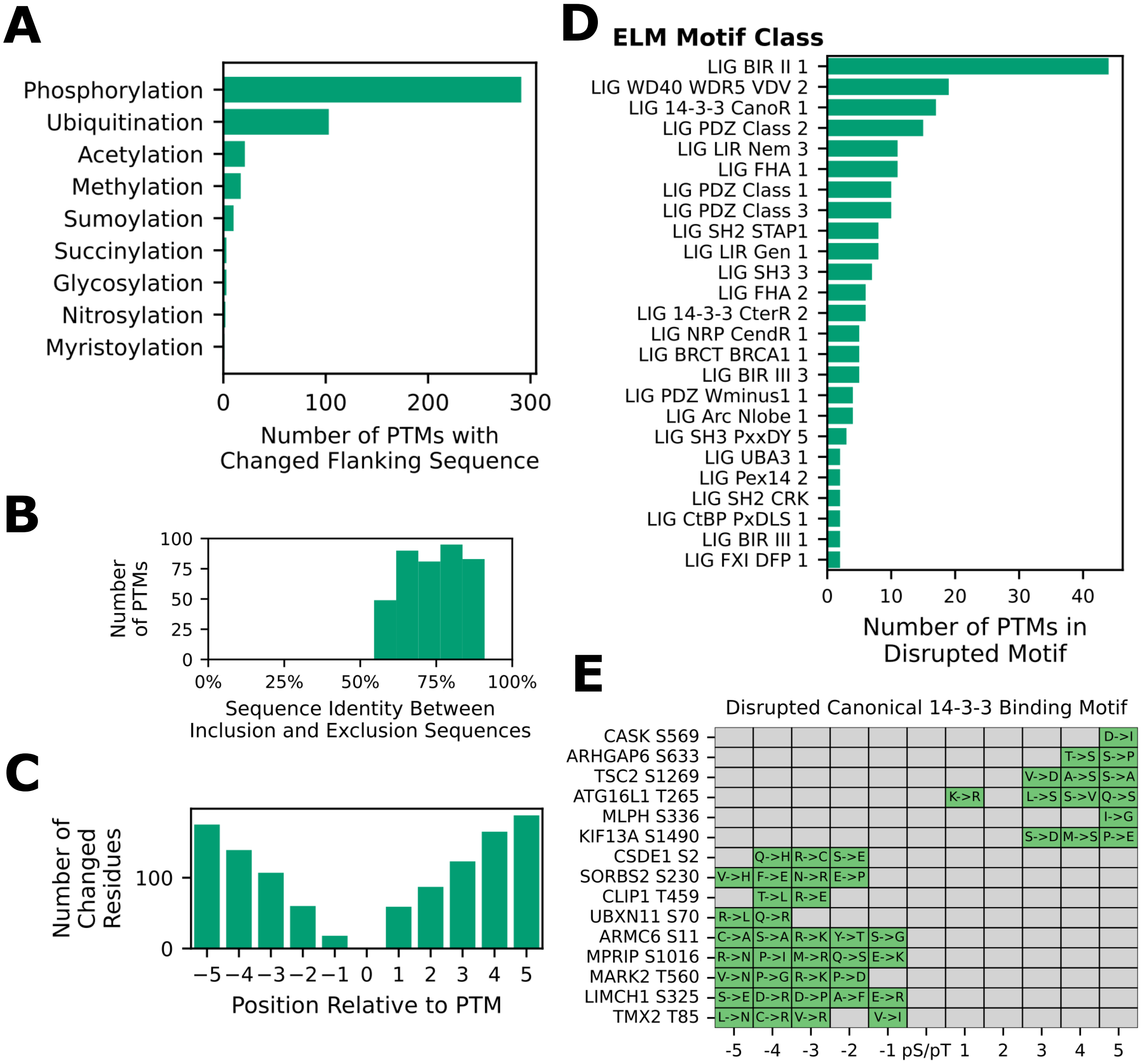


**Supplementary Figure 18. PTMs with altered flanking sequences due to ESRP1 expression changes have the potential to impact binding motifs of interaction domains.** In addition to changes to inclusion of PTMs across isoforms, we identified PTMs nearby splice boundaries whose flanking sequence (five amino acids on either side of modification) are altered with changing ESRP1 expression across TCGA prostate cancer patients. **A)** Number of PTMs of each modification type that are altered as a result of ESRP1 expression, sorted by the most impacted to the least. Only modifications that have altered flanking sequences are shown. **B)** Sequence identity between the different flanking sequences that occur depending on whether the adjacent exonic region is included or excluded. **C)** Positions within the five residue flanking sequence that are most commonly disrupted by changes to ESRP1 expression. The 0 position indicates the modification site. **D)** Number of instances in which a specific ligand binding motif annotated in the Eukaryotic Linear Motif (ELM) database [29] are disrupted, where the ligand motif can be found in the PTM flanking sequence in either the inclusion or exclusion isoform, but not both. Among these motifs are several domains that specifically bind to phosphorylated tyrosines (SH2) or phosphorylated serine/threonines (14-3-3). **E)** Cases in which the canonical ligand binding motif of 14-3-3 proteins is disrupted as a result of ESRP1 expression differences, and the specific residue changes (in green boxes) that occur when going from the inclusion isoform to the exclusion isoform. The motif was identified by the regular expression: 'R[D̂E]{0,2}[D̂EPG]([ST])(([FWYLMV].)—([P̂RIKGN]P)—([P̂RIKGN].{2,4}[VILMFWYP]))'

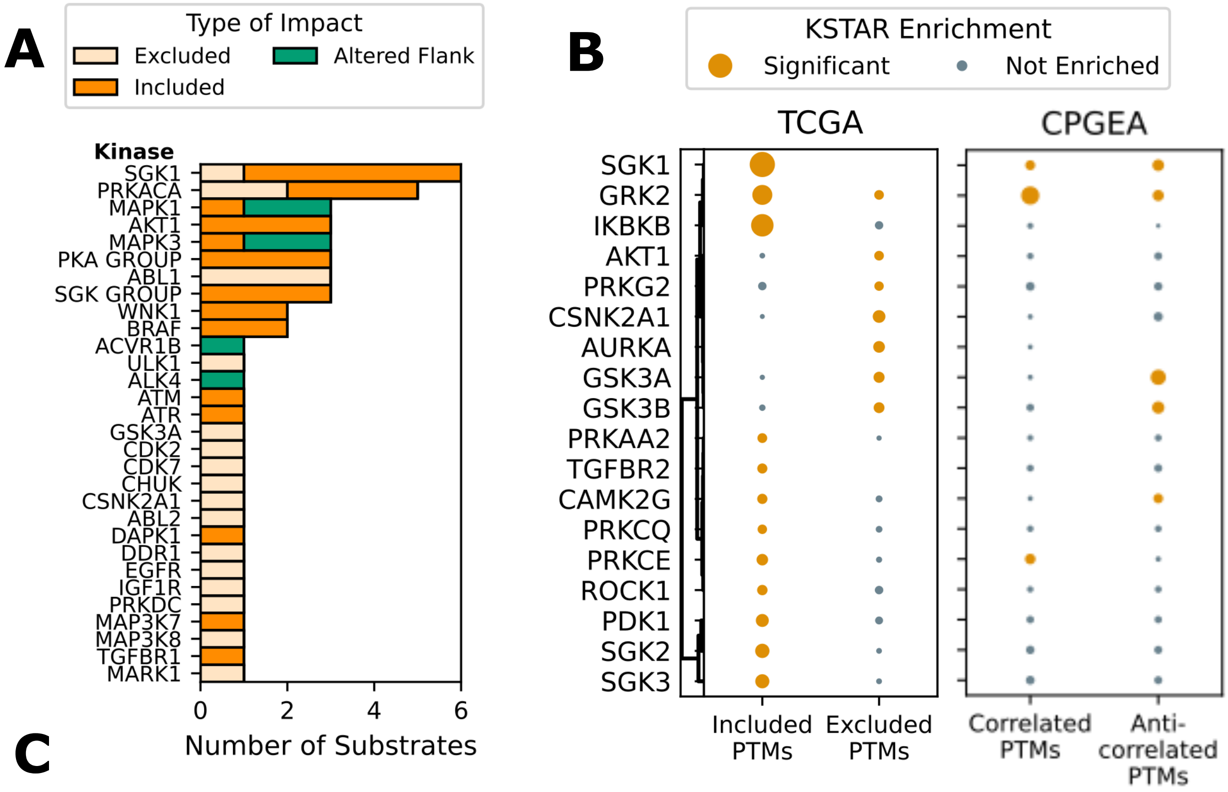


**C**

| Kinase | Site |
|---|---|
| ATP11C | S1126 |
| FLNA | S1981, T2179, T2336 |
| FNIP1 | S232 |
| NEDD4L | S225, S233, S297, S300, T302, S303, S308, S312, T318, S321, S327, S328, S335, S336, S340, **S342**, T344, S365, S366, **T367**, T369, T375, S377, T383, T410, S426, **S428**, S446, **S448**, S449, T451, T453, S455, S464, T473, S475, S479, S483, S487, S538, T540, S795 |
| NSMF | S288 |
| SLTM | S343 |
| TSC2 | **S981** |

**Supplementary Figure 19. ESRP1-regulated splicing increases the availability of SGK substrates.** We sought to explore whether either AKT or SGK exhibited enrichment of available substrates as result of ESRP1-induced splicing changes, which might indicate coordinated changes around a specific kinase/pathway. **A)** Number of kinase substrates among ESRP1-regulated phosphoryation sites, based on known substrates of each kinase annotated in PhosphoSitePlus [4] and RegPhos [13]. Each bar indicates the proportion of a kinase's substrates that are excluded with ESRP1 expression, included with ESRP1 expression, or that exhibit altered flanking sequences. **B)** Median enrichment of predicted substrates for various kinases within ESRP1-related sites identified in the TCGA or CPGEA prostate cancer cohorts. Analysis is based on a modified version of the kinase activity prediction algorithm, KSTAR [30]. Substrate enrichment was assessed for all 50 KSTAR kinase-substrate networks using a hypergeometric test, and the median p-value for each kinase across all 50 networks was extracted. The dotplot shows the median -log10(p), with statistically significant enrichment indicated in orange. All kinases with significant enrichment in either the included or excluded group are shown. **C)** Predicted and known substrates of SGK1 that are increased in ESRP1-high prostate cancer patients. Known substrates are shown in red, the remaining substrates were found in at least 3 of the 50 kinase-substrate networks from KSTAR.

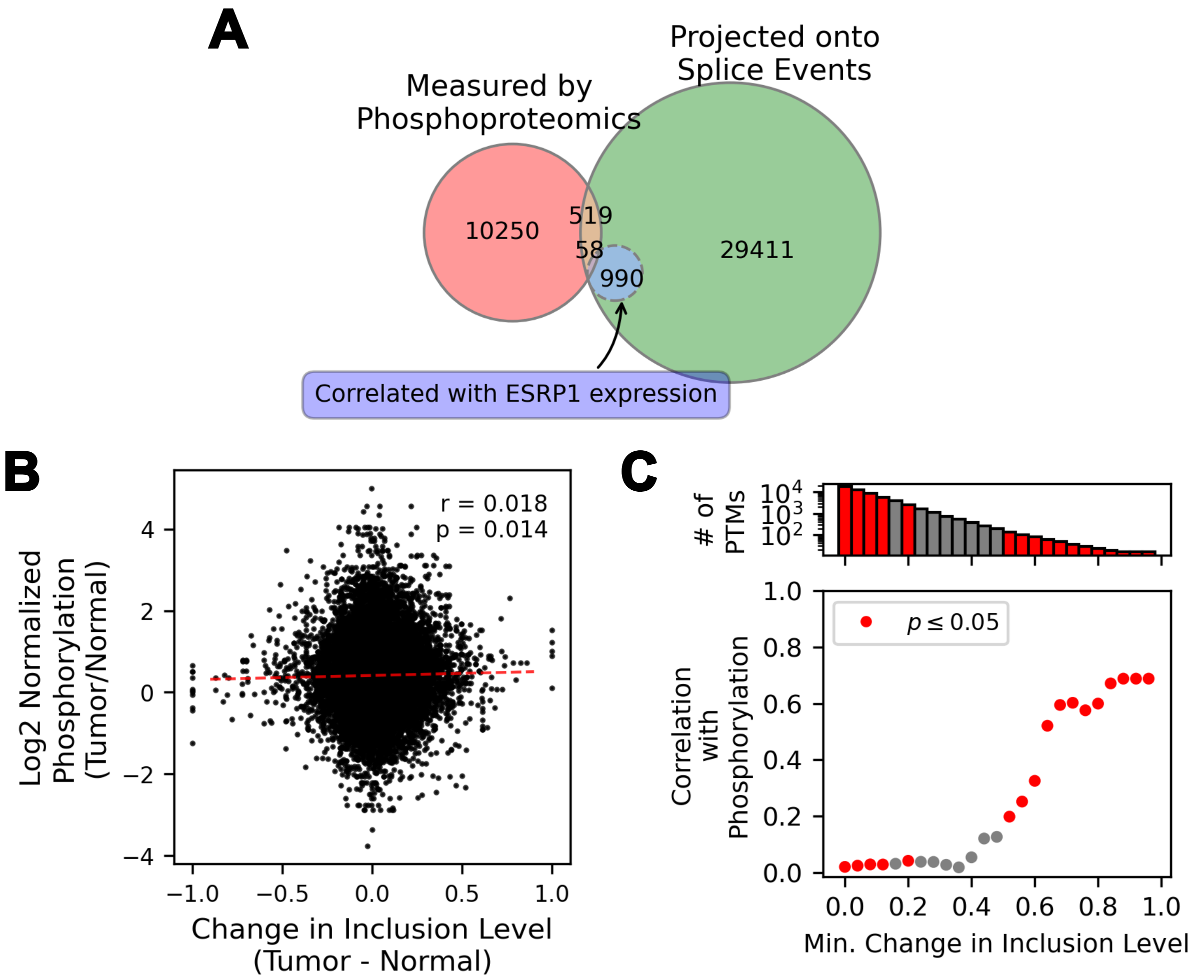


**Supplementary Figure 20. Comparison of differential inclusion of phosphorylatio sites and measured phosphorylation abundance in CPGEA prostate cancer patients** To determine whether we might expect spliced PTMs in prostate cancer patients to lead to changes at the protein level, we applied rMATS [17] and PTM-POSE to an additional prostate cancer cohort from the CPGEA project with matched phosphoproteomic data for each patient [20] [31]. We then compared inclusion of phosphorylation sites to their measured abundance. **A)** Overlap of phosphorylation sites measured by mass spectrometry in the CPGEA prostate cancer cohort and those with splicing quantification available, including those found to be correlated with ESRP1 expression. Only splice events with sufficient variation in inclusion values across samples were analyzed ($range \geq 0.4$, $standard\ deviation \geq 0.05$) **B)** Comparison of the difference in PTM inclusion level between matched normal and tumor tissues and the corresponding measured phosphorylation abundance for sites measured in the CPGEA cohort. **C)** Pearson correlation between inclusion level changes and phosphorylation abundance when removing sites below a minimum change in inclusion level between normal and tumor samples. Significant correlations are noted in red.

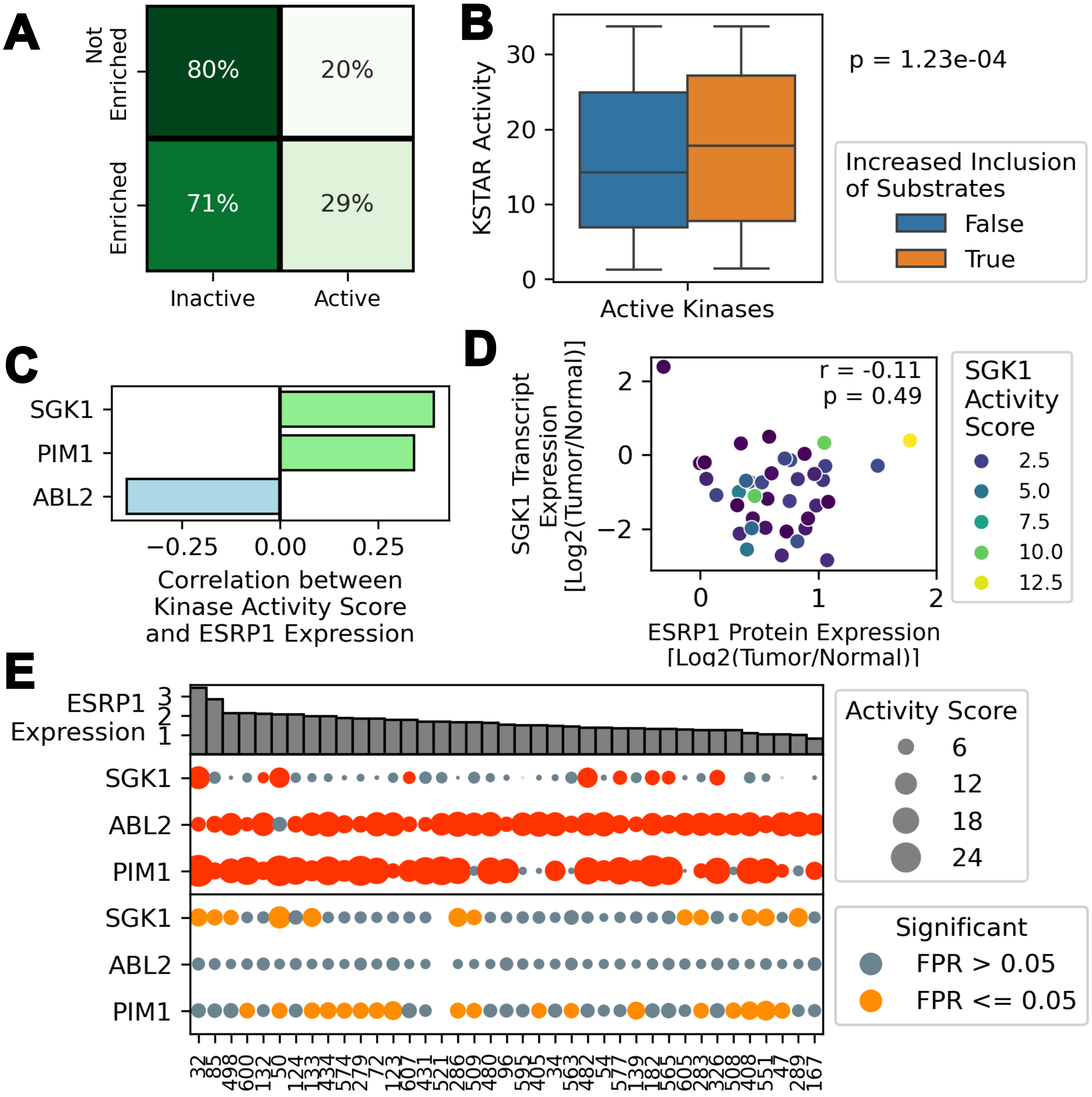


**Supplementary Figure 21. Splicing of phosphorylation sites enhances kinase activity in CPGEA prostate cancer patients.** Using phosphoproteomic measurements for individual tumor samples from CPGEA (relative to matched healthy tissue), we applied a kinase activity inference algorithm, KSTAR [30], to predict kinase activity in each patient. We compared activity predictions to enrichment of a kinases' substrates among phosphorylation sites with increased inclusion ($\Delta PSI \geq 20\%$). **A)** Fraction of cases where kinases with differentially included substrates were also found to be active. **B)** Kinase activity scores for kinases with significant activity within a patient if inclusion of their substrates was also significantly increased due to splicing. Kinases were found to have significantly higher activity scores if splicing led to an increase in their substrates (t-test: $p < 0.05$). **C)** Pearson correlation of SGK1, PIM1, and ABL2 activities with ESRP1 expression across all CPGEA patients. Only kinases with significant correlations are shown. **D)** Relationship between SGK1 FPKM from RNA-sequencing and ESRP1 protein expression in individual CPGEA patients, and its impact on predicted SGK1 kinase activity (indicated by color of each point). Pearson correlation and significance is indicated in upper right of plot. **E)** Kinase activity scores (red) and substrate enrichment among differentially included sites (orange) sorted by ESRP1 expression in CPGEA patients. Size of each dot indicates the degree of activity/enrichment, while colored dots (red/orange) indicate significance ($FPR \leq 0.05$).
